# The endocannabinoid 2-arachidonoylglycerol is released and transported on demand via extracellular microvesicles

**DOI:** 10.1101/2024.09.23.614520

**Authors:** Verena M. Straub, Benjamin Barti, Sebastian T. Tandar, A. Floor Stevens, Tom van der Wel, Na Zhu, Joel Rüegger, Cas van der Horst, Laura H. Heitman, Yulong Li, Nephi Stella, J. G. Coen van Hasselt, István Katona, Mario van der Stelt

## Abstract

While it is known that endocannabinoids (eCB) modulate multiple neuronal functions, the molecular mechanism governing their release and transport remains elusive. Here, we propose an “*on-demand release*” model, wherein the formation of microvesicles, a specific group of extracellular vesicles (EVs) containing the eCB, 2-arachidonoylglycerol (2-AG), is the rate-limiting step. A co-culture model system that combines a reporter cell line expressing the fluorescent eCB sensor, GRAB_eCB2.0_, and neuronal cells revealed that neurons release EVs containing 2-AG, but not anandamide, in a stimulus-dependent process regulated by PKC, DAGL, Arf6, and which was sensitive to inhibitors of eCB facilitated diffusion. A vesicle contained approximately 2000 2-AG molecules. Accordingly, hippocampal eCB-mediated synaptic plasticity was modulated by Arf6 and transport inhibitors. This “*on demand release*” model, supported by mathematical analysis, offers a cohesive framework for understanding eCB signaling at the molecular level and suggests that microvesicles carrying signaling lipids regulate neuronal functions in parallel to canonical synaptic vesicles.

## Introduction

Traditional forms of neurotransmission involve the storage of polar neurotransmitters in synaptic vesicles that fuse with the plasma membrane upon neuronal depolarization, releasing set amounts of neurotransmitters in the synaptic cleft (1). Recently, lipids have emerged as a distinct signaling mechanism to regulate neuronal functions (2). It has been proposed that such lipid messengers, such as the eCBs, anandamide and 2-AG, greatly differ from neurotransmitters as they are not stored in vesicles due to their lipophilic nature. Instead, they are produced by select stimuli and at a specific time and subcellular site (3, 4). This “on-demand production” model is widely used to explain the regulatory role of eCB on neuronal functions (5–7).

2-AG acts as a retrograde messenger produced by the postsynaptic neuron upon depolarization or activation of metabotropic receptors (8, 9). It traverses the synapse and activates the presynaptic cannabinoid 1 receptors (CB_1_R), thereby modulating neurotransmitter release, synaptic plasticity, and neuronal phenotype, as well as complex behaviors related to brain development, learning, memory, appetite, energy balance, pain sensation, and emotional states (10–12). Diacylglycerol lipase-α (DAGLα), a biosynthetic enzyme of 2-AG, plays a crucial role in its “on-demand production” (8, 9). This lipase is highly expressed by neurons and contains a structural motif that binds to Homer proteins, an important component of the molecular scaffold that allows metabotropic glutamate receptor signaling (13). Furthermore, DAGLα activity is regulated by various kinases (14, 15). Genetic deletion or pharmacological inhibition of DAGLα drastically reduces 2-AG levels in the CNS, impairing synaptic plasticity (9), inducing hypophagia (16) and heightening anxiety and fear responses (17, 18).

The termination of 2-AG signaling is hypothesized to occur via its selective uptake from the synaptic cleft by a transport protein, facilitating its diffusion across the plasma membrane (19). To date, the identity of the eCB transport protein remains unknown (20). Once taken up, 2-AG undergoes rapid metabolism by specific enzymes, such as monoacylglycerol lipase (MAGL) and, to a lesser extent, α,β-hydrolase domain-containing protein 6 and 12 (ABHD6 and ABHD12), which together control 2-AG levels in the brain (21). For instance, inhibition of MAGL in the CNS leads to a tenfold increase in 2-AG levels (22), enhancing depolarization-induced suppression of inhibition (DSI) (23), inducing anti-nociceptive behavior (24), and exerting anxiolytic effects (25).

While the current “on-demand production” model captures many features of eCB signaling, it falls short in providing a complete molecular understanding of its release and transport. For example, it fails to explain how hydrophobic lipid messengers like 2-AG (LogP = 6.7) are released from neurons; how the specificity of eCB actions on different cell types is generated; and how they travel from dendrites to CB_1_Rs expressed on axon terminals (3, 26). Hence, a deeper understanding of the molecular mechanism governing eCB release and transport remains to be established.

Studying the dynamic changes in eCB levels, however, has been challenging due to the lack of assays capable of tracking endogenously produced eCBs in a spatial, quantitative, and time-resolved manner. Liquid chromatography coupled to mass spectrometry (LC-MS) and the use of radiolabeled eCB have been pivotal techniques for measuring their levels in the context of their transport (27). Although they are powerful tools for specific quantification, they represent snapshots and fail to capture the spatiotemporal nature of eCB signaling. Recently, a genetically encoded fluorescent sensor, GRAB_eCB2.0_, has been developed that allows spatiotemporal resolved imaging of the dynamic changes in eCB levels produced by neurons in culture, acute brain slices, and brain of freely moving animals (28–32). The sensor has, however, not yet been widely used to address the central question of how eCB are released from neuronal cells.

Here, we propose a new “on-demand release” model, wherein formation of extracellular microvesicles (EVs) containing 2-AG is the rate-limiting step. Using a novel eCB transport assay in which neuronal cells are co-cultured with a cell line expressing the eCB sensor, GRAB_eCB2.0_, we demonstrate that eCBs are released and transported via the emission of EVs containing 2-AG, and not anandamide, in a stimulus-dependent process regulated by protein kinase C (PKC), DAGL, and ADP ribosylation factor 6 (Arf6). Each microvesicle contains approximately 2000 molecules of 2-AG. In addition, we demonstrated that DSI, the prototypical form of eCB-mediated synaptic transmission in acute brain slices, was also modulated by Arf6 and eCB transport inhibitors. Our model, supported by mathematical analysis, provides a comprehensive framework for eCB signaling at the molecular level, potentially explaining why there are different types of endocannabinoids. Our study also suggests an important role for extracellular microvesicles carrying eCB in neuronal communication in the CNS, which works in conjunction with classical synaptic vesicles containing polar neurotransmitters.

## Results

### A GRAB_eCB2.0_-based assay to study eCB release and transport

To explore the molecular mechanism underlying paracrine eCB release, we developed a two-culture system using GRAB_eCB2.0_-expressing HEK293T cells alongside Neuro2A cells in microscopy dishes (Figure 1A). Live cell fluorescence confocal microscopy showed that when Neuro2A cells were stimulated with 1 mM of ATP, which activates P2X_7_ receptors (P2X_7_Rs) and increases 2-AG levels (33), the GRAB_eCB2.0_ signal on the plasma membrane of nearby HEK293T cells were activated and exhibited fluorescence increase. The fluorescence intensity reached a maximum after 3 minutes and then gradually declined (Figure 1B). In contrast, no fluorescence response was detected in co-cultures with HEK293T cells expressing mutant-GRAB_eCB2.0_, which does not respond to high levels of eCBs (Figure 1B).

**Figure 1.**
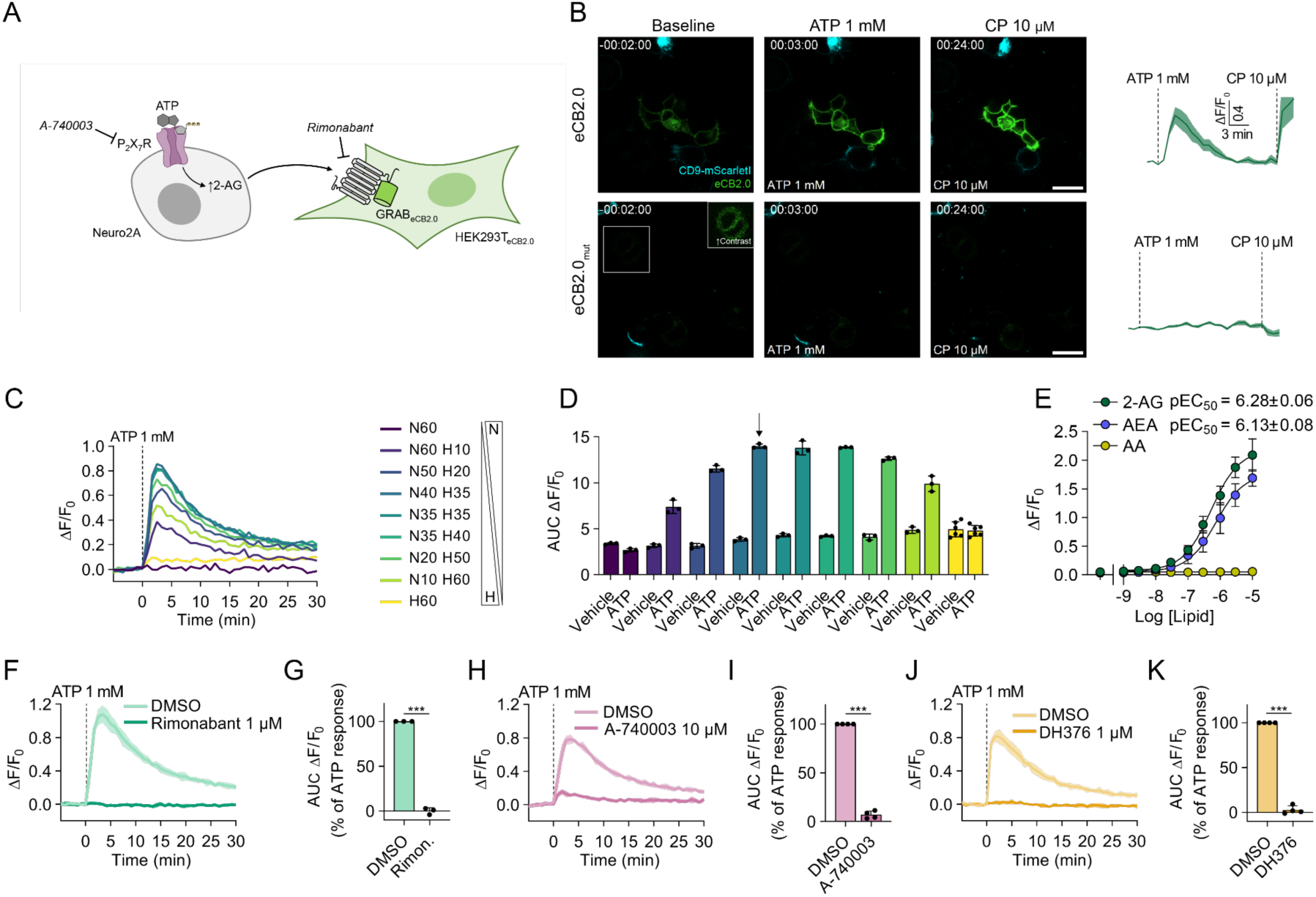
GRAB_eCB2.0_-based co-culture assay to study transcellular 2-AG signaling. **(A)** Schematic representation of the transcellular endocannabinoid transport assay. HEK293T cells expressing GRAB_eCB2.0_ are paired with wild-type Neuro2A cells. Activation of purinergic P2X_7_ receptors (P2X_7_R) on Neuro2A cells by exogenous ATP triggers 2-arachidonoyl glycerol (2-AG) production and release. 2-AG released from Neuro2A cells is free to travel and activate GRAB_eCB2.0_ on HEK293T cells. **(B)** Representative confocal images and traces of HEK293T cells transiently expressing GRAB_eCB2.0_ or GRAB_eCB2.0mut_ and Neuro2A cells transiently expressing CD9-mScarletI. Cells were treated with 1 mM ATP. After 20 min, 10 µM CB_1_ agonist (-)CP-55,940 was added. Traces show mean ΔF/F_0_±SEM of 3-5 ROIs from 2 individual dishes. Scale bars are 20 µm. **(C)** Traces of different number of HEK293T_eCB2.0_ (H) and Neuro2A (N) cells after treatment with vehicle or 1 mM ATP. **(D)** Area under the curve (AUC) of traces after ATP- or vehicle treatment. Arrow indicates optimal ratio (1:0.875, 40.000 Neuro2A + 35.000 HEK293T_eCB2.0_) with a maximum response to ATP and minimal background signal. Data show mean AUC±SD of 3 wells in a 96-well plate. **(E)** Dose-response of GRAB_eCB2.0_ activation in the optimized transport assay by 2-AG, anandamide (AEA) and arachidonic acid (AA). Data are mean±SD, pEC_50_ values are mean±SEM (N=3, n=6). **(F,H,J)** Representative traces (mean±SD; N=1, n=6). Cells were treated with the indicated compounds for 30 min at 37°C prior to baseline measurement. **(G,I,K)** AUC of fluorescent changes, calculated as percentage of vehicle-corrected ATP-response. Data are shown as mean±SD (N=3-4, n=6). Statistical analysis was performed using two-tailed t-test. *** *P* < 0.001.

Next, we transferred the assay to a 96-well format. Using a fluorescence plate reader, we found that the ratio of 35k HEK293T_eCB2.0_ and 40k Neuro2A cells produced the largest response with minimal background (Figure 1C,D). The ATP-induced GRAB_eCB2.0_ signal was effectively blocked by the CB_1_R antagonist, rimonabant (Figure 1F,G) and the P2X_7_R antagonist, A-740003 (Figure 1H,I; Supplementary Figure 2). Treatment with ATP negligible affected the signal of GRAB_eCB2.0_ HEK293Tcells cultured alone (Figure 1C,D), while cannabinoid agonist, CP-55,940, added as a positive control, induced a substantial increase in fluorescence (Supplementary Figure 1C,D).

We also demonstrated the sensor’s performance in the optimized transport assay. Direct application of 2-AG, anandamide and arachidonic acid (AA) showed that 2-AG was the most potent signaling lipid with a pEC_50_±SEM of 6.28±0.06, followed by anandamide (pEC_50_ = 6.13±0.08) (Figure 1E). AA did not activate GRAB_eCB2.0_ up to 10 µM, the highest concentration tested. To verify that the ATP-stimulated fluorescence response was mediated by 2-AG, we applied the potent dual DAGLα/β inhibitor, DH376 (1 µM), which completely abolished the fluorescence response of the sensor (Figure 1J,K). As expected, the ABHD6 inhibitor KT182 did not modulate the fluorescent signal (Supplementary Figure 3A) (33). Together, these findings demonstrate that the two-culture system is suitable to study trans-cellular signaling of endogenously produced 2-AG.

### PKC regulates 2-AG signaling

First, we leveraged our model system to profile an array of small-molecule inhibitors to uncover regulators of 2-AG synthesis, release and transport. Protein kinase C (PKC) has been shown to regulate cycling of DAGL between the plasma membrane and EEA1- and Rab5-positive endosomal compartments via a clathrin-independent pathway (34). Inhibiting PKC reduced endocytosis of DAGLα, thereby increasing the pool of this enzyme at the plasma membrane. In line with this result, we found that the PKC inhibitor Sotrastaurin at 1 µM increased GRAB_eCB2.0_ activation to 127±11%, whereas the PKC activator phorbol 12-myristate 13-acetate (PMA) at 100 nM decreased the GRAB_eCB2.0_ sensor response to 54±7.4% (Figure 2A,B). Notably, PKC is also activated by DAG, the substrate of DAGLα (34), suggesting a potential negative feedback mechanism whereby active PKC reduces DAGLα localization at the plasma membrane, thereby decreasing 2-AG signaling.

**Figure 2.**
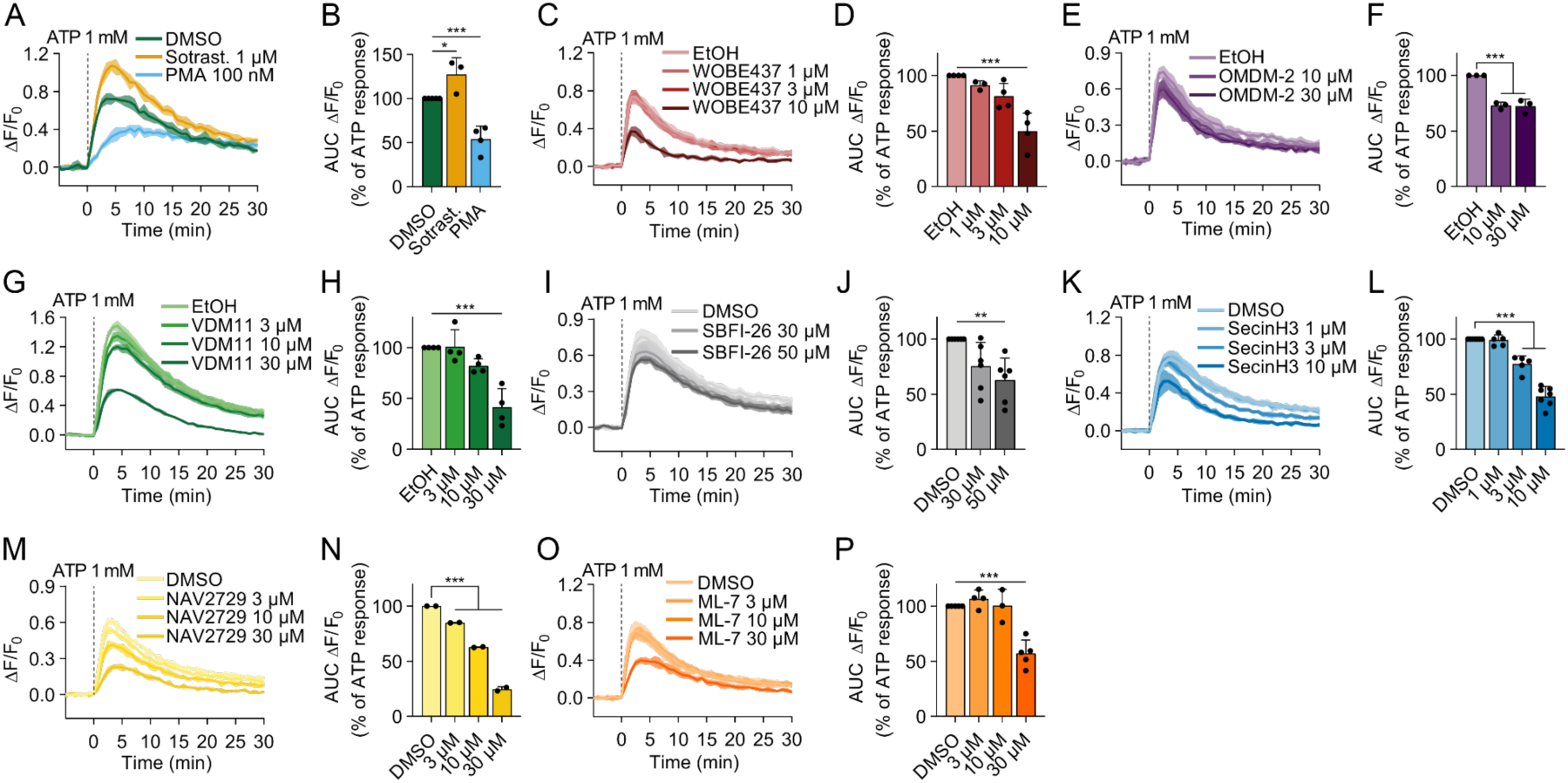
Pharmacological screening reveals regulators of 2-AG release and transport. **(A,C,E,G,I,K,M,O)** Representative traces (mean±SD; N=1, n=6) of changes in fluorescence upon ATP-stimulation in the endocannabinoid transport assay. Cells were treated with the indicated compounds for 30 min at 37°C prior to baseline measurement. **(B,D,F,H,J,L,N,P)** Area under the curve (AUC) of fluorescent changes (ΔF/F_0_), calculated as percentage of vehicle-corrected ATP-response. Data is shown as mean±SD (N=2-7, n=6). Statistical analysis was performed using one-way ANOVA with Tukey’s correction for multiple comparisons. * *P* < 0.05, ** *P* < 0.01, *** *P* < 0.001.

### eCB transport inhibitors reduce 2-AG signaling

Next, we investigated whether the release and transport of 2-AG was mediated via facilitated diffusion, given that eCBs cross the plasma membrane bidirectionally via an unidentified transporter (35). To this end, we tested three structurally different eCB transport inhibitors: WOBE-437 (27), OMDM-2 (36) and VDM-11 (37), which have been shown to inhibit this process. All three inhibitors significantly reduced ATP-induced GRAB_eCB2.0_ signal in a concentration-dependent manner (Figure 2C-H). WOBE437 was the most potent inhibitor and reduced sensor activation by 50±8% at 10 µM, followed by VDM11 and OMDM-2 with 59±9 and 28±4% reduction, respectively, both at 30 µM.

Fatty acid binding proteins (FABPs), especially FABP5, have also been shown to facilitate intracellular, and possibly extracellular, transport of 2-AG (38, 39) and to modulate eCB-mediated synaptic plasticity (40, 41). In our model system, the FABP5 inhibitor, SBFI-26 (42), partly reduced the ATP-induced sensor signal (Figure 2I,J). Taken together, these results suggest that both the eCB transporter and FABP5 may be involved in the release and transport of 2-AG, rather than the reuptake of 2-AG by the Neuro2A or HEK293T cells.

### Inhibitors of extracellular microvesicles release reduce 2-AG signaling

ATP is known to induce EV release in a P2X_7_R-dependent manner (43). Since EVs have been reported as carriers of eCB in N9 microglial cells, primary microglia, and midbrain dopaminergic neurons (44–46), we investigated the role of EVs in the transport of 2-AG. We tested several compounds reported to block different pathways for EV release. Manumycin-A and GW4869, inhibitors of exosome biogenesis via the ESCRT-dependent and ESCRT-independent pathway, respectively (47), did not affect the ATP induced increase in GRAB_eCB2.0_ signal (Supplementary Figure 3C-F). In contrast, SecinH3 and NAV-2729, which primarily target microvesicle release via inhibition of GTPase Arf6 function (48, 49), dose-dependently blocked the ATP induced increase in GRAB_eCB2.0_ signal (Figure 2K-M). The response was reduced to 48±3% and 25±2%, respectively (Figure 2L,N). ML-7, an inhibitor of myosin light chain kinase (MLCK), a downstream effector of Arf6 (49), reduced the ATP induced increase in GRAB_eCB2.0_ signal to 57±6% (Figure 2O,P) (50). Of note, none of the agents significantly affected GRAB_eCB2.0_ signal in HEK293T cells stimulated with CP-55,940, nor was any cytotoxicity observed (Supplementary Figure 4A-J). Together, these data suggested that microvesicle release, but not exosome release, which is controlled by Arf6 and MLCK, is involved in ATP-stimulated 2-AG release and transport.

Finally, we tested whether 2-AG production and release can be induced via activation of the metabotropic bradykinin receptor B_2_ with bradykinin (BK) (51–53) and through a similar mechanisms as activation of the ionotropic P2X_7_R. Indeed, BK treatment of Neuro2a cells led to an increase in GRAB_eCB2.0_ signal on HEK293T cells, which was blocked by DH376, WOBE437 and SecinH3 (Supplementary Figure 5), suggesting a common mechanism for metabotropic and ionotropic-induced 2-AG release and transport.

### Microvesicles carry 2-AG

To obtain an independent line of evidence supporting 2-AG release via extracellular microvesicles, we isolated and characterized EVs from Neuro2A cell culture supernatant using previously reported methods (Figure 3A). ATP increased the EV release by 2-fold as assessed by EV marker protein CD9 detected by western blot, which could be partly blocked by the P2X_7_R antagonist A-740003 (Figure 3B,C). Nanoparticle tracking analysis (NTA) confirmed the increase in number of EVs following addition of ATP (Figure 3D,E). The EVs were of medium to large size ranging from 100 to 600 nm (Figure 3D) with a mean size of 296±19.0 nm (Figure 3G). Stimulation with ATP did not alter the size distribution of isolated vesicles (Figure 3D,F,G). Following ATP stimulation, cells released 7.2±2.4 10^8^ particles, which equates to around 65±25 vesicles released per cell.

**Figure 3.**
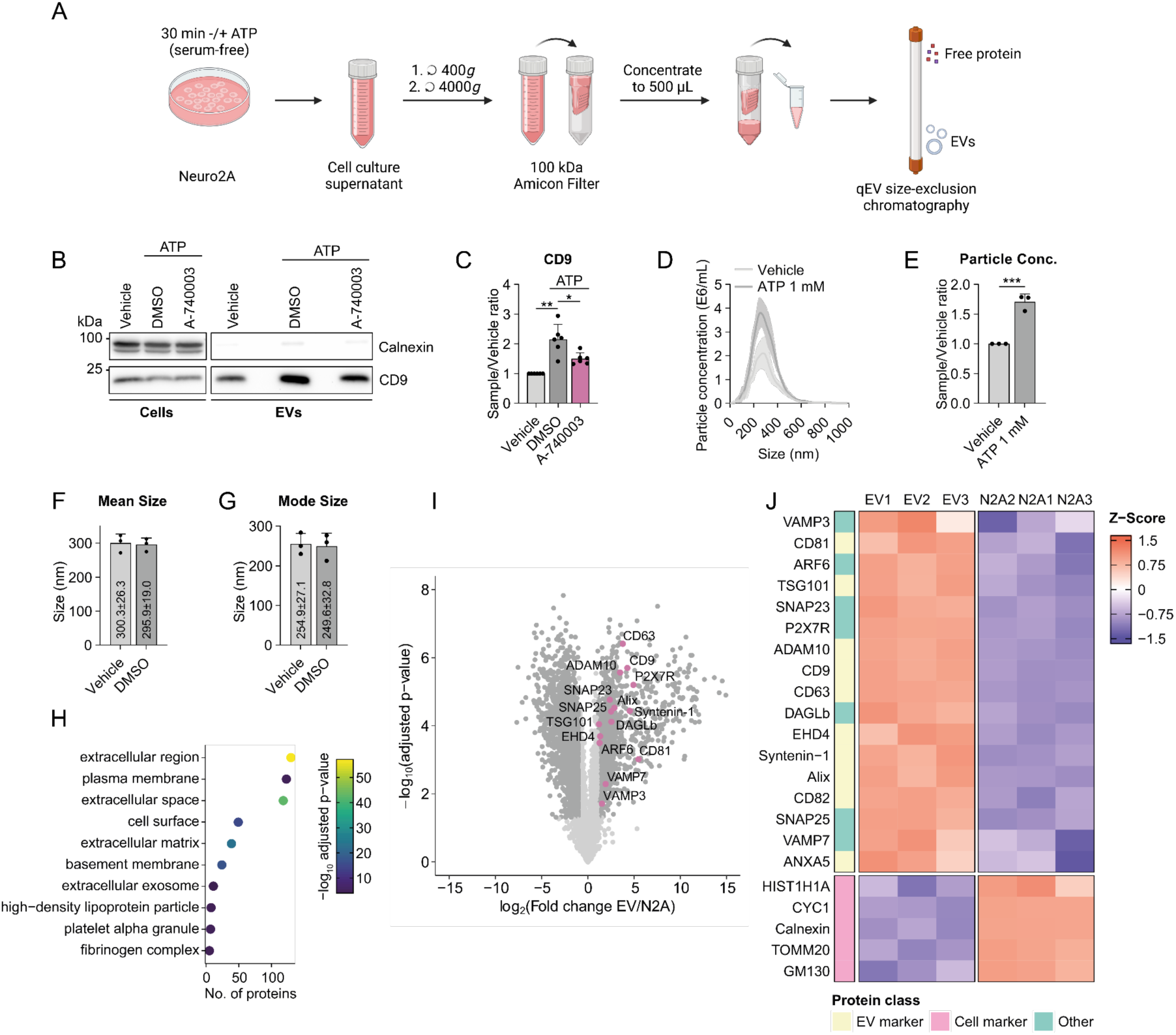
ATP stimulates P2X_7_R-dependent microvesicle release from Neuro2A cells. **(A)** Schematic representation of EV isolation process. Release of EVs is stimulated by 1 mM ATP for 30 min under serum-free conditions. Cell culture supernatant is collected and cleared from cells and cell debris. Supernatant is concentrated to 500 µL using 100 kDa centrifugal filters. EVs are separated from free protein by size-exclusion chromatography (SEC) using Izon qEV 30nm columns. **(B)** Representative western blot of cell lysates and EVs following treatment with DMSO or 10 µM A-740003 (20 min, 37°C) and stimulation with 1 mM ATP or MQ (30 min, 37°C). **(C)** Quantification of CD9 signal intensity. Fold change is calculated relative to the respective vehicle control. Data is shown as mean±SD (N=6). Statistical analysis was performed using matched one-way ANOVA with Tukey’s correction for multiple comparisons. **(D)** Representative size distribution of EVs determined by nanoparticle tracking analysis. Data is mean±SD (N=1, n=3 videos). **(E-G)** Fold change of particle concentration (E), mode size (F), and mean size (G) of EVs as determined by NTA. Data is shown as mean±SD (N=3, n=3 videos). Statistical analysis in (E) was performed using two-tailed t-test. **(H)** Gene ontology enrichment analysis for cellular compartment of proteins enriched >10-fold in EVs compared to Neuro2A cell lysate as determined by LC-MS/MS based proteomics. **(I)** 5181 proteins were identified in EVs. Fold change of protein intensity in EVs compared to Neuro2A cell lysate is shown. Proteins of interest are highlighted. **(J)** Scaled protein abundance of selected proteins in EVs and Neuro2A cells (N2A). EV1-3 and N2A1-3 represent individual biological replicates. * *P* < 0.05, ** *P* < 0.01, *** *P* < 0.001.

We further characterized the EVs by LC-MS/MS based proteomics, which led to the identification of 1160 proteins more than 10-fold enriched in EV preparations compared to Neuro2A cell lysate. Proteins related to the extracellular space/region, plasma membrane, and extracellular exosome were overrepresented by cellular component gene ontology analysis (Figure 3H). Multiple EV marker proteins were detected, including CD9, CD63, CD81, CD82, ADAM10, SNARE proteins, syntenin-1, EHD4 and Annexin-A5 (Figure 3I,J). In turn, marker proteins for several cellular organelles (HIST1H1A, CYC1, Calnexin, TOMM20 and GM130) were significantly reduced in EVs compared to Neuro2A cell lysates, validating the identity and purity of the isolated EVs (Figure 3J). Of note, ARF6, FABP5 and SCP-2 were also found in the EV fractions. Further analysis of molecular functions and biological pathways associated with proteins enriched in EVs revealed that integrin binding, receptor binding, heparin binding, and collagen binding were significantly overrepresented in EVs compared to the cell lysate. Among the biological pathways, cell adhesion and cell-matrix adhesion were two of the three most enriched (Supplementary Figure 6). Taken together, this suggests that these vesicles contain several components of the 2-AG release and transport machinery and may be targeted to the surface of specific recipient cells.

Next, we probed whether eCB and related lipids were contained in the EVs using LC-MS/MS based targeted lipidomics. We measured a lipid panel of 26 signaling lipids, comprised of 2-AG, anandamide and their analogues as well as free fatty acids. Out of the 26 lipid species, 17 could be identified above detection levels in Neuro2A EVs (Supplementary Figure 7), including 2-AG and AA, but not anandamide. In addition to 2-AG, monoacylglycerols 2-oleoylglycerol (2-OG) and 2-lineoyl glycerol (2-LG) were identified. Of the free fatty acids, oleic acid, linoleic acid, palmitic acid, stearic acid, eicosatrienoic acid (ETA) and dihomo-*γ*-linolenic acid (DGLA) were found in EVs. Additionally, *N*-stearoylethanolamine, *N*-palmitoylethanolamine, *N*-oleoylethanolamine, *N*-linoleoyl ethanolamine and *N-*pentadecanoyl ethanolamine were detected in EVs.

Remarkably, 2-AG and AA levels were significantly increased to 139±7% and 153±13%, respectively, in the EV preparations following ATP stimulation (Figure 4A). Importantly, 2-AG was not detected in protein-containing fractions 9+10 (data not shown) and cellular levels of all lipids remained unchanged (Figure 4B, Supplementary Figure 8). ETA, DGLA, 2-OG and 2-LG levels in EVs were slightly increased, but these changes did not reach statistical significance (Supplementary Figure 7). In contrast to the studies of Gabrielli et al. (44) and Lombardi et al. (45), we could not detect anandamide in Neuro2A EVs, which could be due to differences in the EV isolation protocol or, more likely, due to the different cell types studied.

**Figure 4.**
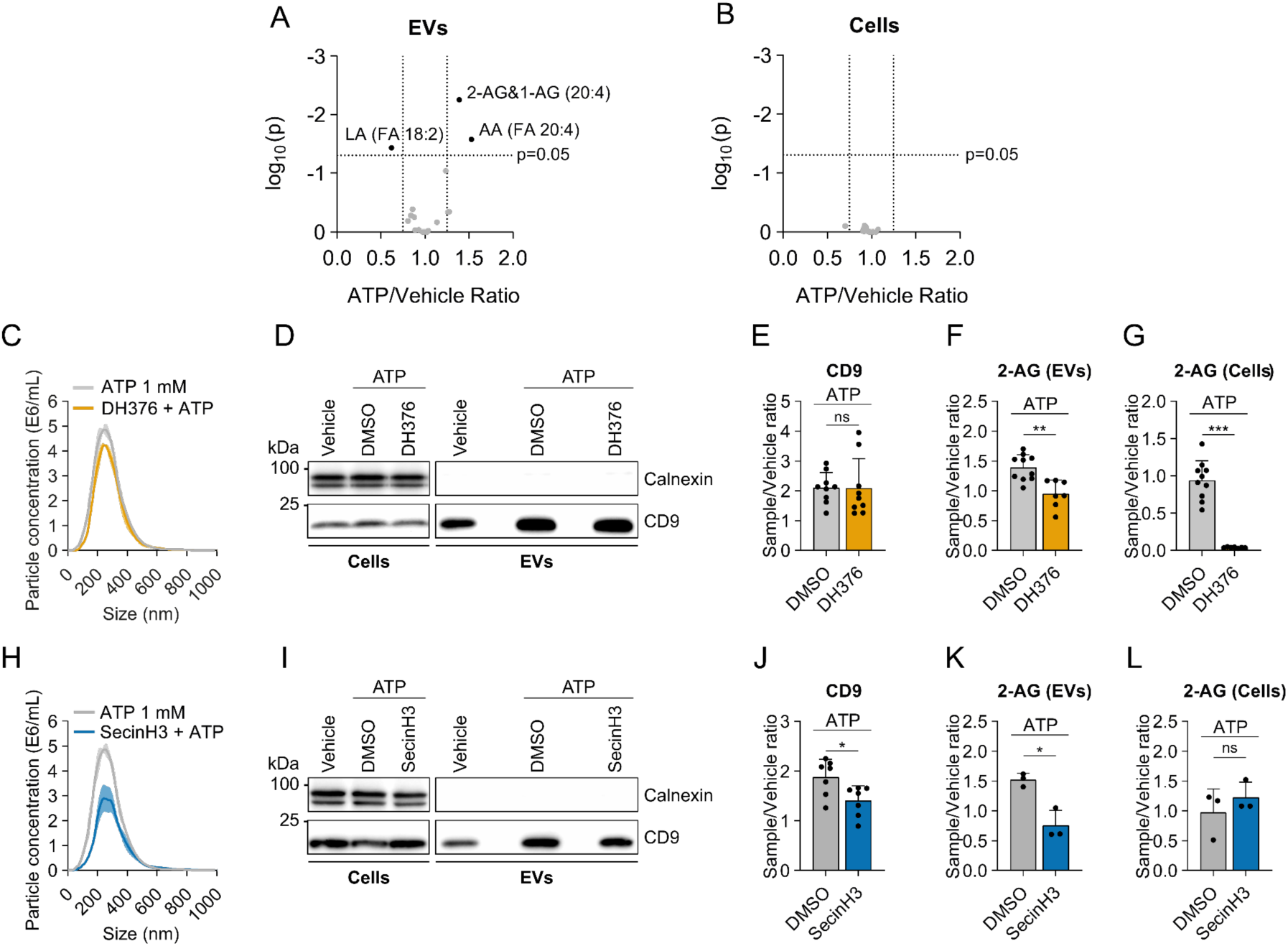
2-AG is specifically sorted into microvesicles in a DAGL- and Arf6-dependent process. **(A,B)** Fold-enrichment (ATP/Vehicle) of lipids in EVs (A) or cells (B) following vehicle (MQ) or ATP-treatment for 30 min at 37°C. **(C,H)** Size distribution of EVs determined by nanoparticle tracking analysis. Data are mean±SD (N=1, n=3 videos). **(D,I)** Representative western blot of cell lysates and EVs. **(E,J)** Quantification of CD9 signal intensity. Fold change is calculated relative to the respective vehicle control. **(F,G,K,L)** 2-AG levels in EVs (F,K) and cells (G,L) relative to vehicle-treated control. Cells were treated with 1 µM DH376 (C-G) or 10 µM SecinH3 (H-L) for 20 min prior to vehicle (MQ) or ATP-treatment for 30 min at 37°C, followed by EV isolation. Data in E-G and J-L are shown as mean±SD (N=3-10 EV isolations). Statistical testing was performed using one-way ANOVA with Tukey’s correction for multiple comparisons. * *P* < 0.05, ** *P* < 0.01, *** *P* < 0.001, ns not significant *P* > 0.05.

Taking the average particle count of EVs released by ATP-stimulated cells as determined by NTA (7.2±2.4×10^8^) and the average absolute 2-AG levels (2507±706 fmol), each vesicle contained an estimated number of 2000 2-AG molecules. If all isolated vesicles contain 2-AG to the same extent, the average number of 2-AG molecules per vesicles with or without ATP stimulus remained constant. This would suggest that the increased number of vesicles released by ATP is sufficient to explain the GRAB_eCB2.0_ response. However, 2-AG and AA were the only lipids significantly increased in the total pool, therefore it is conceivable that ATP triggers the release of specific EV subsets that are enriched in 2-AG and AA levels. In summary, we show that ATP induces the release of extracellular microvesicles from Neuro2A cells that contain 2-AG, but not anandamide.

### DAGL and Arf6 regulate release of 2-AG in microvesicles

Subsequently, we investigated whether the ATP-induced increase in 2-AG in EVs depends on DAGL activity. Incubation of Neuro2A cells with 1 µM of DH376 prior to ATP-stimulation and EV isolation did not alter the number or size-distribution of released EVs (Figure 4C-E), but the increase of 2-AG and AA content in EVs was blocked (Figure 4F, Supplementary Figure 7). This suggests that 2-AG produced by DAGL is loaded into EVs and that its incorporation can be uncoupled from the release of EVs. In turn, we next asked whether a reduction in EV release influences the 2-AG content detected in EVs. To this end, we interfered with ATP-stimulated EV release by SecinH3-mediated inhibition of Arf6 activity. Western blot analysis showed a 43±19% reduction in CD9 levels in line with the particle analysis (Figure 4I,J). 2-AG levels in cells were not changed following SecinH3 treatment (Figure 4L). In EV preparations, however, SecinH3 blocked the increased levels of 2-AG following ATP stimulation (Figure 4K). This showed that while SecinH3 does not influence 2-AG biosynthesis itself (because cellular 2-AG levels remained unchanged), it reduced the number of EVs and thereby the total content of 2-AG in the extracellular space. Of note, the 2-AG level in the EVs was only 0,25% of the total cellular pool. The small extracellular pool compared to the bulk 2-AG cellular levels, which were both dependent on DAGL biosynthesis, indicate that 2-AG remains in the cell and does not spontaneously diffuse into the aqueous extracellular environment. Taken together, this suggests that 2-AG is specifically sorted into extracellular microvesicles in a Arf6-dependent mechanism and released upon an ATP stimulus.

### Depolarization-induced suppression of inhibition involves Arf6

Next, we aimed to translate our key findings to an acute hippocampal slice model, where the molecular and cellular organization of the brain circuit remains largely intact, to investigate whether Arf6 plays a role in 2-AG-mediated synaptic plasticity. First, we confirmed that primary hippocampal neurons produce 2-AG in a stimulus-dependent manner via DAGL and Arf6, using our transport assay (Supplementary Figure 9). Next, we examined whether depolarization-induced suppression of inhibition (DSI), a prototypical form of eCB-mediated synaptic plasticity, is controlled by Arf6 and eCB transport inhibitors. To test this, we conducted paired-patch clamp recordings between CB_1_ receptor-positive basket cells and pyramidal cells in acute hippocampal slices, infusing Arf6 or eCB transport inhibitors into the postsynaptic neuron via the patch pipette (Figure 5A). We found that the synaptic charge transferred during IPSCs, a strong readout for DSI efficacy, was affected by SecinH3, NAV2729 and WOBE437 (Figure 5B-F). Differences were already apparent shortly after the start of the experiment and became even more pronounced after more than 20 min of recording (Figure 5C-F). Importantly, the synaptic charge remained unaffected during baseline synaptic transmission when DSI was not induced (Figure 5B). In contrast, the inhibitors reduced DSI (Figure 5D-F), indicating a specific alteration of phasic 2-AG-mediated signaling without affecting baseline synaptic activity. Notably, the initial amplitude of DSI was unaffected by the inhibitors, suggesting a delayed onset of the effects of the intracellularly applied inhibitors. Considering the 20 minutes intracellular drug infusion of the postsynaptic cell, the delayed onset is unlikely attributed to technical factors such as slow diffusion. Together, these results suggest that phasic 2-AG signaling in hippocampal slices involves Arf6 and the unidentified endocannabinoid membrane transporter, closely aligning with key findings from our transport assay using cultured cells.

**Figure 5.**
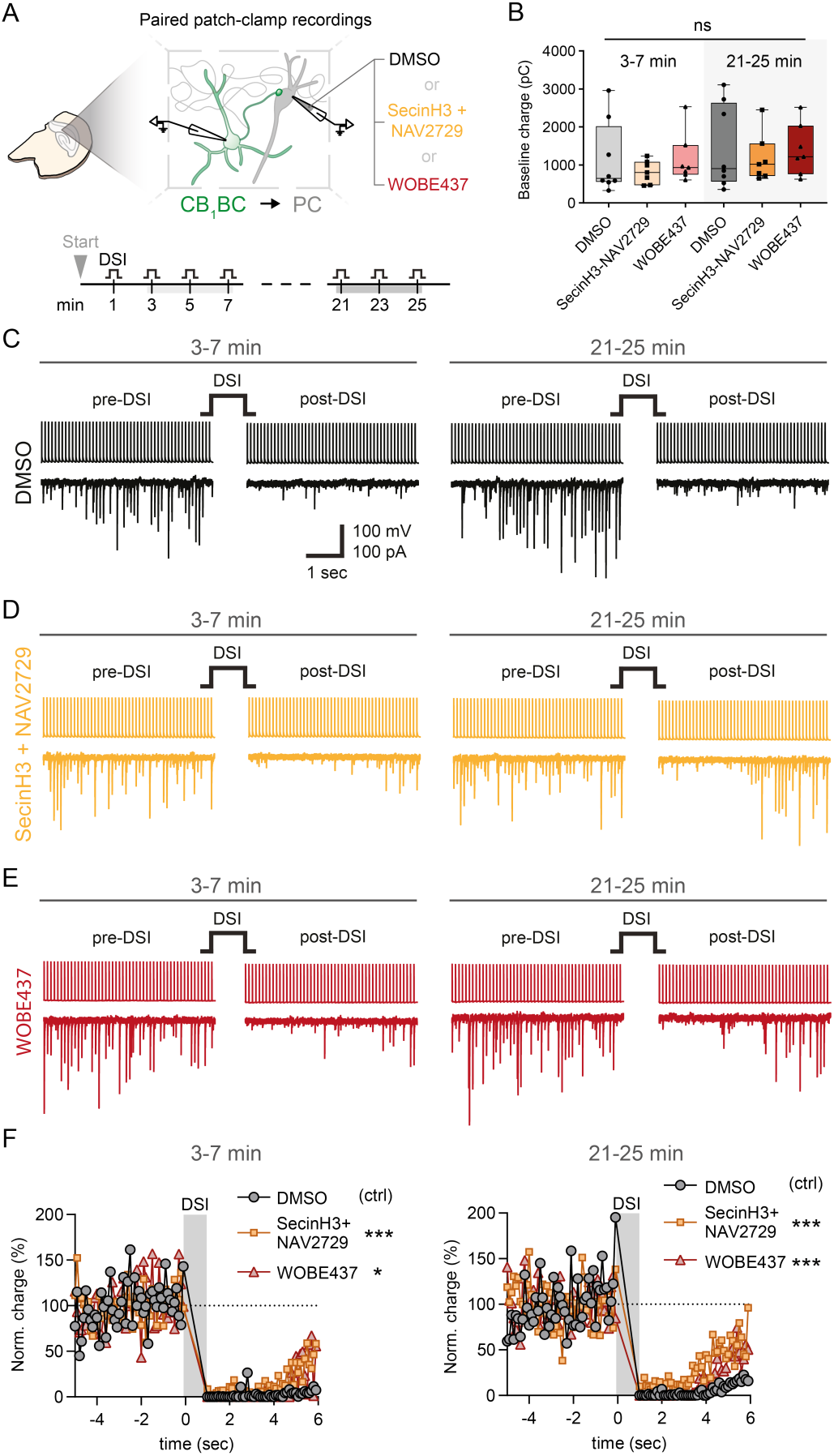
Inhibition Arf6 and facilitated diffusion affects 2-AG dependent phasic endocannabinoid signalling *ex vivo*. **(A)** Schematic illustration of experimental design. Paired patch-clamp recordings were conducted in acute hippocampal slices between CB_1_-positive basket cells (CB_1_BC) and pyramidal cells (PC). The intracellular solution for the postsynaptic cell contained either vehicle DMSO, SecinH3 and NAV2729 or WOBE437 in the pipette. Depolarization induced suppression of inhibition (DSI) were induced with 1 sec depolarization of the postsynaptic cell in every 2 minutes after the start of the experiment. The timeline of the experiment represents analyzed time windows. **(B)** Summary graphs of baseline synaptic charge between pairs. **(C)** Representative traces of presynaptic action potential evoked (top traces) inhibitory postsynaptic currents (IPSCs, bottom traces) before and after DSI induction at 3-7 min and 21-25 min time windows. Postsynaptic intracellular solution contained vehicle 0.5% DMSO. **(D)** Same as C, but postsynaptic intracellular solution contained SecinH3 (10 µM) and NAV2729 (30 µM). (**E)** Same as C-D, but postsynaptic intracellular solution contained WOBE437 (10 µM). **(F)** Summary plots of normalized charge values before and after DSI induction at 3-7 min and 21-25 min time windows. Data show median ± IQR with individual datapoints (B) or mean (F). Statistical significance was determined by Kruskal-Wallis test with Dunn’s multiple comparisons (B) or repeated measures one-way ANOVA with Dunnett’s multiple comparisons (F). * P < 0.05, *** P < 0.001, ns not significant P > 0.05. N= 7 animals / group n = 8 / 7 / 7 individual pairs in groups DMSO / SecinH3 and NAV2729 / WOBE437, respectively

### Formation of extracellular microvesicles containing 2-AG is the rate-limiting step

Finally, to determine the rate-limiting step in 2-AG release, we developed a mathematical model of reporter cell activation (Figure 6A and Supporting Information). The model considers 2-AG production, metabolism and distribution between a main Neuro2A cellular 2-AG pool (m_cell_) and an EV pool located in the plasma membrane (m_EV_). The EV pool represents 2-AG sorting into the forming microvesicles. An ATP stimulus releases EVs containing 2-AG from the EV pool into the extracellular space. A fraction of the extracellular 2-AG is absorbed by the adjacent cells in which 2-AG is distributed and metabolized. The 2-AG concentration in the reporter cell determines the observed fluorescent signal strength generated by the GRAB_eCB2.0_ sensor.

**Figure 6.**
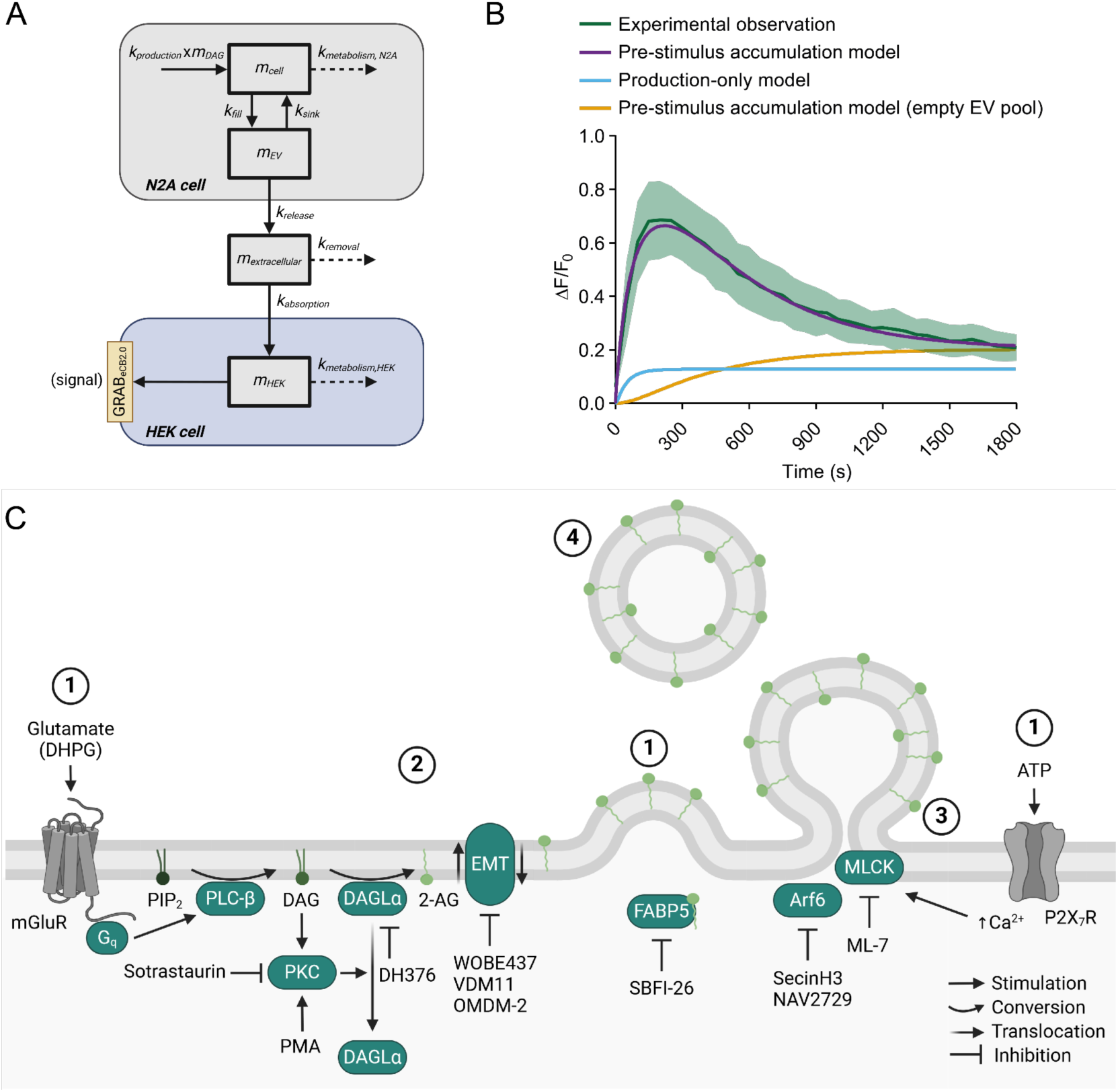
Model for on-demand 2-AG release in extracellular microvesicles. **(A)** Schematic of the mathematical compartmental model to approximate GRAB_eCB2.0_ activation in the endocannabinoid transport assay. **(B)** Model fitting shows that the implementation of a preformed EV pool is important to describe the formation of the observed signal peak. A model that only takes 2-AG production into account was unable to capture the dynamics the signal peak. **(C)** Scheme on the on-demand release model (1) Stimulation of the cell leads to lipid shuffling and cytoskeleton rearrangements, resulting in the formation of a budding plasma membrane microvesicle. (2) 2-AG is loaded into forming microvesicles. This process is regulated by: Production of 2-AG by DAGLα; PKC-dependent endocytosis of DAGLα; Translocation of 2-AG within the lipid bilayer by an unidentified endocannabinoid transporter; and intracellular endocannabinoid transport proteins. (3) Release of budding vesicles through membrane fission is controlled by Arf6- and MLCK-activity. (4) 2-AG containing extracellular vesicles released into the extracellular space mediate cell-to-cell communication and may be taken up by various cell types to terminate endocannabinoid signaling. 2-AG: 2-arachidonoyl glycerol, Arf6: ADP-ribosylation factor 6, ATP: Adenosine triphosphate, DAG: Diacyl glycerol, DAGLα: DAG lipase α, DHPG: (S)-3,5-dihydroxyphenylglycine, EMT: Endocannabinoid membrane transporter, FABP5: Fatty-acid binding protein 5, G_q_: heterotrimeric G protein alpha subunit q, mGluR: metabotropic glutamate receptor, MLCK: Myosin light chain kinase, P2X_7_R: P2X_7_ receptor, PIP_2_: Phosphatidylinositol-4,5-bisphosphate, PKC: Protein kinase C, PLC-β: Phospholipase C-β, PMA: Phorbol 12-myristate 13-acetate.

The model was fitted to describe and quantify the characteristics of the experimentally observed signal dynamics of ATP-stimulated Neuro2A cells (Figure 6B and Supporting Information). Notably, the inclusion of extracellular microvesicle pool in Neuro2A cells in the model was necessary to capture the dynamics of signal peak formation observed at approximately 220 s after the ATP stimulus. In contrast, an alternative model structure which takes only 2-AG production into account, and not the 2-AG sorting into forming microvesicles, could not replicate the formation of the signal peak (Figure 6B and Supporting Information), suggesting the presence of a microvesicle pool in Neuro2A cell before the ATP stimulus.

Using the model, we estimated that 59% of 2-AG molecules in the extracellular space was originally present in the microvesicle pool and released in 38 vesicles per Neuro2A cell during the initial burst. The remaining 2-AG was released in a distinctly lower rate of one vesicle every approximately 67 seconds after the initial signal burst. The fill rate constant k_fill_ of the microvesicle pool was 5.7 × 10^-7^*s*^-1^, which is significantly smaller than the 2-AG production rate constant k_prod_ (6.7 × 10^-3^*s*^-1^). Consequently, the model predicts that it takes substantial time (> 60 min) to completely refill the m_EV_ pool with 2-AG upon depletion. Of note, the absorption rate constant k_abs_ (1.3 × 10^-7^*s*^-1^) was approximately 5-fold lower than k_fill_. In summary, the model suggested that the formation of microvesicles loaded with 2-AG is the rate-limiting step in eCB release from Neuro2A cells, while the uptake of 2-AG by the reporter cell is the overall rate-determining step in GRAB_eCB2.0_ sensor activation.

## Discussion

There are three current hypotheses to explain cellular eCB trafficking: a) (intra)cellular transport via lipid carrier proteins (40, 41); b) transport and release via extracellular vesicles, (44–46) and c) release and reuptake via unidentified membrane transporters (27). Yet, the molecular mechanism how eCBs are released from neuronal cells is poorly understood (11). To address this important subject, we developed an eCB reporter cell line by introducing the eCB sensor GRAB_eCB2.0_ into HEK293T cells (28), and combined it in a 96-well plate format with mouse neuroblastoma Neuro2A cells, commonly used for studying eCB signaling (54). Leveraging this model system, we demonstrated that activated neuronal cells release microvesicles containing 2-AG in a process regulated by PKC, DAGL, Arf6, and which is sensitive towards inhibitors of eCB facilitated diffusion. Key findings were replicated in primary hippocampal neurons and acute hippocampal slice preparations. We demonstrated that DSI, a prototypical form of eCB-mediated synaptic transmission, was also modulated by Arf6 and eCB transport inhibitors. Based on the literature (*vide infra*), our experimental findings and mathematical analysis, we propose an “on-demand release” model, where the formation of extracellular microvesicles containing 2-AG constitutes the rate-limiting step (see Figure 6C). This model involves the following four steps:

1. *Activation of neurons leads to membrane budding.* The activation of neurons induces membrane budding. This process is triggered by stimuli like synaptic activity (43, 55), metabotropic and ionotropic receptor stimulation, leading to calcium-dependent rearrangements of the cytoskeleton (55) and membrane lipids (56–58), culminating in the protrusion of the plasma membrane (58–60).
2. *Sorting of 2-AG into outward budding plasma membrane:* 2-AG is sorted into the outward budding plasma membrane, which relies on its biosynthesis by DAGLα at the plasma membrane (29, 61). DAGLα activity and its subcellular localization is dynamically regulated via kinases like PKA, PKC and CaMKII (14, 15) and interactions with Homer-proteins situated in neuronal subdomains close to the metabotropic glutamate receptor and rich in the EV marker Flot-1 (13). The transportation of 2-AG into outward budding plasma membrane may involve eCB transporters, which could act as lipid scramblases facilitating 2-AG diffusion between bilayer leaflets (62), although further research is needed to fully understand this process. Lipid carrier proteins like FABP5, HSP70, and SCP2 may play an additional role in transporting 2-AG from other organelles, as well as lipid droplets, to the forming microvesicles (40, 41). The cargo-loading of microvesicles also involves SNARE proteins (60, 63, 64), with postsynaptic synucleins and synaptobrevin (VAMP2) being necessary for the release of eCB (31). Thus, the sorting of 2-AG into vesicles is governed by multiple pathways, likely influenced by the cell type, subcellular origin of 2-AG and the specific subset of vesicles released.
3. *Membrane fission leads to microvesicle release*: The connection between newly formed vesicle and the cell membrane is eventually severed through membrane fission. Arf6 plays a crucial role by recruiting myosin light chains kinase (MLCK) to phosphorylate MLC at the vesicle neck, leading to actin cytoskeleton remodeling and EV release (49, 58, 65). Notably, we found a delayed onset of action of the Arf6 and transport inhibitors in the modulation of hippocampal synaptic transmission. This might suggest that there is a pre-existing pool of outward budding vesicles in the plasma membrane filled with 2-AG. Once this pool is released, the subsequent time-dependent formation of new microvesicles and their loading with 2-AG become dependent on Arf6 and the putative eCB transporter.
4. *Removal of extracellular microvesicles limits 2-AG signaling:* EVs containing eCBs like 2-AG play a crucial role in cell-to-cell communication in the central nervous system (CNS) (44–46, 66). Signaling lipids can be efficiently transferred from microvesicles to plasma membranes upon collision or membrane fusion rather than via phagocytosis or endocytosis, which is a slower process (67, 68). The activation of CB_1_R depends on several factors: the number of vesicles interacting with the target cell, the concentration of 2-AG within the vesicle, and the 2-AG diffusion gradient, which is influenced by the distribution of 2-AG and the activity of 2-AG metabolizing enzymes such as MAGL, ABHD6, ABHD12, and MGAT in the CB_1_R-expressing target cell. Additionally, astrocytes and microglia may absorb EVs from the synaptic cleft, potentially limiting 2-AG signalling and contributing to retrograde endocannabinoid signalling at the synapse (69, 70).

### Implications of the on-demand release model

The release of microvesicles containing 2-AG from cells may reflect the metabolic state of a cellular network, as 2-AG is part of the futile lipid cycle characterized by a high metabolic flux, where its biosynthesis and metabolism are in equilibrium (71). Release of 2-AG-containing vesicles enable neighboring cells expressing CB_1_R to sense the network’s metabolic state. In addition, an activity-dependent rapid increase in 2-AG production or microvesicle release, such as due to neuronal activity would then serve to control network activity through CB_1_R-mediated inhibition of neurotransmitter release.

Anandamide was not detected in microvesicles derived from the plasma membrane. The biosynthetic machinery of anandamide is primarily found on presynaptic, intracellular membranes (72), and anandamide can function as an intracellular messenger, thereby acting in an autocrine-like manner (73). This may suggest that the intracellular anandamide concentration reflects activation of presynaptic metabolic processes. Since the signaling pool of 2-AG was only 0.25% of its total cellular pool, this indicates that anandamide and 2-AG concentrations available to activate CB_1_R are in the same order of magnitude. We propose, therefore, that the CB_1_R expressed on presynaptic terminals or astrocytes can act as a sensor integrating the metabolic state from both post- and presynaptic sites by detecting the para- and autocrine signaling of eCBs. Intriguingly, this may also explain why there are two different types of eCB. Anandamide could represent the metabolic state of presynaptic sites, whereas 2-AG reflects post-synaptic metabolic processes in a cellular network.

EVs contain (glyco)proteins, such as lectins and integrins, which may facilitate the targeting of EVs to specific cell types or sites expressing cognate receptors. Interestingly, this could potentially explain the cell-type specificity of eCB signaling (26, 74). Moreover, EVs may extend beyond the synapse, activating CB_1_R at distant sites (*e.g.* on perisomatic GABAergic terminals (3, 75)) or even entering the general circulation, allowing eCB to travel in the body and convey information to cells in other organs about the metabolic state of their site of origin.

To conclude, the development of a two-culture model featuring a genetically encoded fluorescent sensor capable of detecting eCB levels, enabled the time-resolved study of eCB production, release, and trafficking between cells. In combination with electrophysiological studies and mathematical analysis, this led us to propose a new “on-demand release” model, wherein the formation of extracellular microvesicles containing 2-AG is the rate-limiting step. This model reconciliates the three previously proposed hypotheses for eCB signaling. It provides a cohesive and quantitative framework for understanding of eCB signaling at the molecular level that can be tested by future experimental studies. Given that CB_1_R is one of the most abundant G protein-coupled receptor in the brain and eCBs are found in every brain region constituting diverse neuronal circuitries, our study suggests an important role of extracellular microvesicles carrying hydrophobic signaling lipids as a mechanism to regulate neurotransmission in the nervous system in addition to classical synaptic vesicles containing polar neurotransmitters.

## Materials & Methods

### Compounds

SecinH3 (S7685), ML-7 (S8388), GW4896 (S7609) and Sotrastaurin (S2791) were purchased from Selleck Chemicals. A-740003 (HY-50697), and NAV-2729 (HY-112473) were purchased from MedChemExpress. ManumycinA (M6418) and ATP (A2383) were purchased from Sigma-Aldrich. SBFI-26 (AOnareB31865) was purchased from Aobious. PMA (10008014), (-)CP-55,940 (90084), Bradykinin (15539), (S)-3,5-DHPG (14411), 2-arachidonoyl glycerol, anandamide, arachidonic acid, VDM11 (10006731) and OMDM-2 (10179) were purchased from Cayman Chemical. WOBE437 (76) and DH376 (61) were prepared in-house.

### Cell culture

HEK293T, HEK239T_eCB2.0_ and Neuro2A cells (ATCC CCL-131™) were cultured in Dulbecco’s Modified Eagle’s Medium High Glucose (DMEM, Sigma-Aldrich D6546 or Capricorn DMEM-HXA) supplemented with 10% (v/v) heat-inactivated newborn calf serum (Biowest, S0750), 2 mM GlutaMAX™ (Gibco, 35050038) and penicillin and streptomycin (200 µg/mL each, Duchefa Biochemie) at 37°C and 7% CO_2_. Culture medium was replaced after 2-3 days and cells were subcultured twice a week. Cultures were regularly tested for mycoplasma infection and discarded after 3 months.

### Plasmids

For the generation of pPGK_CD9_mScarletI_Flag DNA coding for CD9_Flag was obtained as a gBlock from IDT DNA and cloned into a pcDNA3.1 vector containing a PGK promoter using restriction enzyme cloning. mScarletI with a GGGGS linker was then inserted between the CD9 and the Flag sequence. pcDNA3.1_eCB2.0 and pcDNA3.1_eCB2.0_mut were kind gifts from Nephi Stella. The sequence encoding for GRAB_eCB2.0_ was amplified by PCR and subcloned into a pLenti6.3 backbone using general restriction enzyme cloning methods to generate pLenti6.3_eCB2.0. Identity of sequences was verified by Sanger Sequencing (Macrogen).

### Generation of a stable HEK293T_eCB2.0_ cell line

Virus was produced by transfecting 70% confluent HEK293T cells in a 10 cm dish with pLenti6.3_eCB2.0 and the packaging vectors pMD2.G, pRSV-REV, and pMDLg/pRRE in a 4.17:1.08:1:1.92 (10 µg DNA in total) ratio. After 8 h, medium was replaced with complete DMEM containing 20 mM HEPES. 48h post-transfection, medium was harvested, centrifuged at 3000 g for 10 min and 0.45 µm sterile filtered. 5 mL of virus-containing medium was diluted 1:1 with complete DMEM and added to a 10 cm dish of HEK293T cells for 32 h. Cells were then passaged and cultured in medium containing 10 µg/mL BlasticidinS. After 3 passages the cultures were considered virus-free and cultured as described above with addition of 5 µg/mL BlasticidinS at every other passage.

### Transient transfection

Cells were seeded, grown to 70-80% confluency in 6 cm culture dishes and medium was refreshed with complete DMEM. Plasmid DNA (1 µg/dish) was combined with polyethyleneimine (PEI, 3 µg/dish) in serum-free medium and incubated for 15 min at room temperature. The mixture was slowly added to the cells. After 24 h transfected cells were harvested and seeded for live-cell microscopy, which was performed 48 h post-transfection.

### Live cell-microscopy

Neuro2A cells transfected with pPGK_mScarletI_CD9 and either HEK293T cells transfected with pcDNA3.1_eCB2.0/pcDNA3.1eCB2.0_mut_ or HEK293T_eCB2.0_ cells were washed once with Dulbecco’s phosphate-buffered saline (DPBS, Sigma-Aldrich, D8537) and detached by trypsinization. Cell count and viability were assessed by Trypan Blue Exclusion using a TC20^TM^ automated cell counter (Bio-Rad). 80,000 HEK293T cells and 110,000 Neuro2A cells were seeded in a 35 mm glass-bottom dish (ibidi, 81158). The following day, medium was replaced with 540 µL phenol-red free medium containing 0.1% DMSO or inhibitor and incubated for 20 minutes in the incubator before placing the dish on the microscope stage in a humidified OkoLab chamber at 7% CO_2_, 37°C. Images were acquired with a Nikon Ti2-E inverted microscope equipped with a Nikon A1 confocal head, Galvano scanner, HP Apo TIRF 100xAC/1.49 NA oil immersion objective, 488nm and 561nm lasers, 405/488/561/647nm dichroic mirror and 525/50 and 595/50 filter cubes. The automated correction collar was adjusted to the recommended setting for 37°C. The microscope was controlled using Nikon NIS Elements Software. An area of interest was centered with HEK293T cells and Neuro2A cells in close proximity. Focus was adjusted so that the center of the HEK cells were in focus and Nikon Perfect Focus System was enabled to keep cells in focus throughout the acquisition. Channels were acquired sequentially (total exposure time 32s), with an image acquired every minute. Three images with 1024×1024 pixel size were acquired as a baseline, then 60 µL of 10x ATP 1 mM/vehicle was added directly to the dish and gently but swiftly mixed by pipetting, and 20 more images were collected.

Analysis of images was performed in ImageJ/FIJI. Regions of interests were manually drawn around individual groups of cells and the mean intensity was measured. Change in fluorescence was calculated in relation to the average baseline intensity:

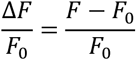

Area under the curve (AUC) was calculated in GraphPad Prism 8.3/9.0. Baseline value was manually set based on the lowest ΔF/F_0_ value in the dataset.

### Plate reader based GRAB assay

Inner wells of a clear-bottom black 96-well plate (Greiner, 655090 or Thermo Scientific, 165305) were coated with 50 µL of 50 µg/mL poly-L-lysine (Sigma-Aldrich, P2636) in 1x borate buffer pH 8.8 (Thermo Scientific, 28341) for 20 min and washed with 100 µL sterile water twice. Outer wells were filled with 100 µL DPBS (Sigma-Aldrich, D8537). Neuro2A cells and HEK293T_eCB2.0_ cells were harvested by trypsinization and counted by Trypan Blue Exclusion using a TC20^TM^ automated cell counter (Bio-Rad). Neuro2A (4E5 cells/mL) and HEK293T_eCB2.0_ (3.5E5 cells/mL) cells were mixed and 100 µL of cell suspension was added per well (40.000 Neuro2A + 35.000 HEK293T_eCB2.0_). The following day, medium was gently aspirated and replaced with 95 µL of pre-warmed Hank’s balanced salt solution (HBSS, Gibco^TM^, 14025092 or Sigma-Aldrich, H8264). 5 µL of 20x compound or DMSO was added (final DMSO conc. 0.1% (v/v)) and cells were incubated at 37°C for 30 min. Plate was placed in a microplate reader (CLARIOstar®, BMG LABTECH), focal height was adjusted based on 2-3 individual wells, and gain was adjusted based on the full plate (target value 10%). A baseline measurement of 7 cycles was taken (bottom read, spiral scan, 3 mm, 30 flashes, 50s cycle time, 37°C, excitation 470/15, emission 515/20). Plate was removed and 5.26 µL of 20x ATP (final conc. 1 mM), Bradykinin (final conc. 10 µM), or vehicle (MQ) in HBSS was swiftly added. Plate was placed back and measured for 37 cycles (30 min). ΔF/F_0_ and AUC were calculated as described above for live-cell microscopy experiments.

Assays for WOBE437, OMDM2, VDM11 were performed in HBSS containing 0.1% (w/v) fatty-acid free BSA (Sigma-Aldrich, A8806) and the vehicle for these compounds was ethanol (final conc. 0.1% (v/v)). Final DMSO concentration for GW4896 assays was 0.3% (v/v). Buffer for NAV2729 assays was HBSS (0.1% BSA (w/v)) and final DMSO concentration was 0.3% (v/v). For EC_50_ determination of lipids, 95 µL of HBSS containing 0.1% (w/v) fatty-acid free BSA was added and the plate was incubated for 30 min. After the baseline measurement (see above), 5 µL of 20x lipid or vehicle was added and the response was measured for 10 cycles. EC_50_ values were determined based on maximum ΔF/F_0_ values for each lipid (average of last 4 cycles). EC_50_ curves were fitted in GraphPad Prism 8.3/9.0.

### Effect of (-)CP-55,940 activated GRAB_eCB2.0_

60.000 HEK293T_eCB2.0_ cells were seeded per well as described above. The following day, medium was gently aspirated and replaced with 95 µL of pre-warmed Hank’s balanced salt solution (HBSS, Gibco^TM^, 14025092 or Sigma-Aldrich, H8264) and cells were equilibrated for 30 min at 37°C. A baseline measurement was taken (see above), 5 µL of 20x (-)CP-55,940 (final conc. 10 µM,) was added, and 7 more cycles were measured. Then, 5.26 µL of 20x compound or vehicle was added and fluorescence was recorded for 30 min.

### LDH Cytotoxicity assay

Cytotoxicity of compounds was tested using the CyQUANT™ LDH Cytotoxicity Assay (Invitrogen). Neuro2A and HEK293T_eCB2.0_ cells were harvested by pipetting and seeded in 96-well plates (Sarstedt, 83.3924) coated with 250 µg/mL poly-L-lysine (10.000 Neuro2A + 8.750 HEK293T_eB2.0_ cells per well). The following day, medium was aspirated and replaced with 95 µL of pre-warmed Hank’s balanced salt solution (HBSS, Gibco^TM^, 14025092 or Sigma-Aldrich, H8264). 5 µL of sterile water was added to control cells and 5 µL of 20x compound/DMSO was added to test wells. Cells were incubated for 1 h at 37°C. 50 µL of cell culture supernatant was transferred to a new plate and LDH activity was measured according to the manufacturers’ instructions.

### EV isolation

Neuro2A cells were seeded in two 15 cm dishes per sample in phenol red-free medium and grown to ∼70 % confluency on the day of harvesting. Cells were gently washed with DPBS once and 20 mL serum-free phenol red-free medium was added. 833 µL of 20x ATP (final concentration 1 mM) or MilliQ (vehicle) was added to the medium, mixed by gently swirling the dish, and incubated for 30 min at 37°C, 7% CO_2_. Cell culture medium of two dishes was combined and centrifuged first at 400*g* for 20 min, then at 4000*g* for 30 min. The supernatant was transferred to a new tube and either used immediately for EV purification or stored at 4°C over night. Cells were harvested by scraping in 2 mL DPBS and counted by Trypan Blue exclusion. Samples were only included if cell viability was >95%. Aliquots of 1 mL of cell suspension were pelleted at 1000*g* for 5 min, snap-frozen in liquid nitrogen, and stored at -80°C.

15 mL 100 kDa Amicon filters (Merck) were washed once with MilliQ and once with particle-free DPBS (Sigma-Aldrich, D1408, diluted to 1x with MilliQ) by adding 5 mL to the filter and centrifuging at 2000*g* for 2 min. Conditioned cell culture medium was added to the filter and was centrifuged at 2000-3000*g* in 2 min intervals until concentrated to below 400 µL. Concentrated medium was retrieved and residual sample was collected by washing the filter with 100-200 µL of DPBS. If necessary, the sample was brought to 500 µL with DPBS.

qEV 35nm original legacy columns (Izon) were used according to the instruction manual in combination with an Automated Fraction Collector (Izon, software v2.2.8).The void volume was set to default. The column was washed with 1.5 column volumes of DPBS (15 mL) before applying 500 µL of sample to the top of the column. Once the sample has entered the column, DPBS was added and 500 µL fractions were collected. EV containing fractions (1&2) were pooled and 30 µL was set aside for Western Blot analysis before storing EVs at -80°C. For NTA, EVs were stored at 4°C and either analyzed directly following isolation or the next day.

### Nanoparticle tracking analysis (NTA)

Particle count and size distribution were analyzed using a Nanosight LM20 (Malvern Pananalytical) instrument. Sample chamber was thoroughly rinsed with MilliQ before first use and between samples. In case of protein build-up the sample chamber was cleaned with 100% isopropanol. Sample was introduced using a syringe pump. Flow rate was initially set to 150 µL/min. Once a stable flow was achieved flow was reduced to 15 µL/min. The laser was focused using 200 nm Nanosphere^TM^ size standards (3200A, ThermoScientific^TM^). EV samples were injected without prior dilution. Camera level was set to 16 and three 90s videos were acquired for each sample. Analysis of videos was performed using Nanosight NTA 2.3 software. Detection level was set to 7 Multi, blur to 3×3, and minimum track length and minimal expected particle size were set to automatic.

### Cell lysis

Cell pellets were thawed on ice and lysed by osmotic shock with a HEPES/sucrose lysis buffer (20 mM HEPES pH 7.2, 1 mM MgCl_2_, 250 mM sucrose, 2.5 U/mL Benzonase^®^ (Santa-Cruz sc-202391)) on ice for 30 min with occasional vortexing. Protein concentration was determined using Qubit Protein Assay (Invitrogen). Lysates were diluted to the desired concentration with 20 mM HEPES pH 7.2, flash-frozen in liquid-nitrogen, and stored at -80°C.

### Immunoblot

10 µL of 4x non-reducing Laemmli buffer (240 mM Tris, 8% (w/v) SDS, 40% (v/v) glycerol, 0.04% bromophenol blue) was added to 30 µL of EVs or 0.5 mg/mL cell lysate and heated to 60°C for 10 min. 26 µL was loaded on a 12.5 % SDS-PAGE gel alongside PageRuler™ Plus Prestained Protein Ladder (ThermoScientific^TM^), leaving an empty slot between ladder and sample to avoid reduction of proteins. Proteins were resolved for 75 min at 180 V. Proteins were transferred to polyvinylidene difluoride membranes using a Trans-Blot Turbo Transfer System (Bio-Rad). Membranes were washed once with TBS (50 mM Tris pH 7.5, 150 mM NaCl), once with TBS-T (50 mM Tris, 150 mM NaCl, 0.05% Tween 20), and blocked with 5% milk in TBS-T for 1 h at RT. Membranes were decorated with primary antibody (rat anti-CD9, eBioscience, 14-0091, 1:1000; rabbit anti-calnexin, Sigma-Aldrich, C4731, 1:3000) diluted in 5% milk/TBS-T at 4°C over night, washed three times with TBS-T, and incubated with HRP-conjugated secondary antibody (goat anti-rat IgG, Cell Signaling, 7077, 1:3000; mouse anti-rabbit IgG, Santa-Cruz, sc-2357, 1:5000) in 5% milk/TBS-T for 1 h at RT. Membranes were washed three times with TBS-T, once with TBS, and incubated with Luminol solution (250 µg/mL Luminol, 100 mM Tris pH 8.8, 67 µM *p*-coumaric acid, 0.3 % H_2_O_2_) for 90s. Chemiluminescence was detected with a ChemiDoc MP System (Bio-Rad).

### Lipid Extraction

800 µL of the combined EV or protein fractions was transferred to a safe-lock 2 mL tube and concentrated to below 400 µL in a vacuum concentrator (30°C, 3h). Neuro2A cell pellets were thawed on ice. Lipid extraction was performed on ice. To each sample, 10 µL of internal standard mix (2-arachidonoyl glycerol-d_8_, *N*-arachidonoyl ethanolamine-d_8_, *N*-docosahexaenoyl ethanolamine-d_4_, *N*-linoleoyl ethanolamine-d_4_, *N*-oleoyl ethanolamine-d_4_, *N*-palmitoyl ethanolamine-d_5_, *N*-stearoyl ethanolamine-d_3_, N-eicosapentaenoyl ethanolamine-d_4_ and arachidonic acid-d_8_, Cayman Chemical Company) was added and samples were vortexed. 100 µL of 0.5% (w/v) NaCl solution was added followed by 100 µL of 100 mM NH_4_CH_3_CO_2_, pH 4. Then, 1 mL of methyl tert-butyl ether was added, and samples were thoroughly mixed using a Next Advance Bullet Blender (7 min, speed 8) before centrifugation at 25 000*g*, 4°C for 11 min. 925 µL of the upper organic layer was transferred to a new tube and evaporated in an Eppendorf Concentrator until dry (1h, V-AQ, 30°C). Lipids were reconstituted in 30 µL (MeCN:H_2_O 9:1) and 25 µL was transferred to LC-MS vials with inserts.

### LC-MS/MS analysis of lipids

10 µL of the reconstituted lipids was measured using an Acquity UPLC I class binary solvent manager pump in conjugation with a tandem quadrupole mass spectrometer as mass analyzer (Waters Corporation, Milford, USA). The separation was performed with an Acquity HSS T3 column (2.1 × 100 mm, 1.8 μm) maintained at 45°C. The aqueous mobile phase A consisted of 2 mM ammonium formate and 10 mM formic acid, and the organic mobile phase B was acetonitrile. The flow rate was set to 0.55 mL/min; initial gradient conditions were 55% B held for 0.5 min and linearly ramped to 60% B over 1.5 min. Then the gradient was linearly ramped to 100% over 5 min and held for 2 min; after 10 s the system returned to initial conditions and held 2 min before next injection. Electrospray ionization-MS and a selective Multiple Reaction Mode (sMRM) was used for endocannabinoid quantification. MRM transitions were individually optimized using their synthetic standards for target compounds and internal standards. Peak area integration was performed with MassLynx 4.1 software (Waters Corporation). The obtained peak areas of targets were corrected by appropriate internal standards peak area to calculate response ratios. Response ratios of cell pellets were normalized to cell count.

### Sample preparation for proteomics

EVs were isolated from the supernatant of 6 15 cm dishes per sample as described above. 0.5 mL 10 kDa Amicon filters (Merck) were washed once with MilliQ before the combined EV fractions were added and concentrated to below 100 µL. Concentrated EVs were retrieved and brought to 100 µL with DPBS if necessary. Cell pellets were lysed and protein concentration was determined as described above. 10 µg of cell lysate was diluted to 100 µL with DPBS. Samples were then prepared for proteomic analysis using S-Trap^TM^ micro columns (Protifi^TM^). 100 µL of 2x SDS lysis buffer (10% SDS, 100 mM TEAB, pH 8.5) was added to all samples, vortexed and briefly sonicated using a Qsonica Q700MPX Microplate Horn sonicator (3x 10s). Cell lysates were cleared by centrifugation at 13,000*g* for 8 min and only the supernatant was further processed. 8.7 µL of freshly prepared reductant was added (final conc. 5 mM TCEP) and incubated at 55°C for 15 min. Samples were allowed to cool down for a few minutes before 8.7 µL of alkylator was added (final conc. 20 mM MMTS) and incubated for 10 min at RT. Samples were acidified to below pH 1 by adding 21.7 µL of 27.5% phosphoric acid and vortexing. Then, 1435 µL of binding/wash buffer (100 mM TEAB/90% methanol, pH 7.55) was added and samples were mixed. Samples were transferred to S-Trap micro columns in 150 µL steps, centrifuging at 3,000*g* for 30 s to trap proteins. Columns were washed six times with binding/wash buffer and centrifuged at 3,000*g* for 30 sec. Columns were rotated 180° between centrifugations. After the last wash, residual buffer was removed at 4,000*g* for 1 min. Columns were moved to a clean low-protein binding tube (Eppendorf) and 25 µL digestion buffer was added (1 µg sequencing grade modified trypsin (Promega, V511A) in 50 mM TEAB). S-Traps^TM^ were capped loosely and incubated while stationary at 37°C in a water-saturated atmosphere for 18 h. Peptides were eluted in three rounds by adding 40 µL and centrifuging at 4,000*g* for 1 min. First with 50 mM TEAB, then with 0.2% formic acid in water, and last with 50% acetonitrile. All elutions were combined and peptides were dried in a vacuum concentrator at 45°C for 1-2 h until dry. Samples were reconstituted in 30 µL LC-MS solution (3% (v/v) acetonitrile, 0.1% (v/v) formic acid, 10 fmol/μl yeast enolase internal standard), vortexed, and mixed at 1500 rpm for 5 min. Reconstituted peptides were centrifuged at 20,000*g* for 10 min and peptide concentration in the supernatant was measured by NanoDrop. EV samples were brought to 300 ng/µL with LC-MS solution and then all samples were transferred to LC-MS vials and stored at 4°C.

### MS data acquisition

Peptide samples were analyzed using a nanoElute 2 LC system (Bruker) coupled to a timsTOF HT mass spectrometer (Bruker). 5 µL of sample was loaded on a trap column (PepMap C18, 5 mm x 0.3 mm, 5 µm, 100 Å, Thermo Scientific) followed by elution and separation on the analytical column (PepSep C18, 25 cm x 75 µm, 1.5 µm, 100 Å, Bruker). A gradient of 2 - 25% solvent B (0.1% FA in ACN) in 25 min, 25 - 32% B in 5 min, 32 - 95% in 5 min and 95% B for 10 min at a flow rate of 300 nL/min (all % values are v/v, water, TFA and ACN solvents were purchased from Bisolve, LC-MS grade). ZDV Sprayer 20 µm (Bruker) installed in the nano-electrospray source (CaptiveSpray source, Bruker) was used with following source parameters: 1600 V of capillary voltage, 3.0 L/min of dry gas, and 180°C of dry temperature. To determine the ideal amount of dia-PASEF isolation windows the average peak width was determined in DDA PASEF mode with a full ion mobility window of 0.6 to 1.6 Vs/cm2 and 10 PASEF ramps in a mass range from 100 m/z to 1700 m/z with charge states from 0 to 5+. The dual TIMS analyzer was utilized under a fixed duty cycle, incorporating a 100 ms ramp time, resulting in a total cycle time of 1.17 s. Precursors that reached a target intensity of 20,000 (intensity threshold 2,500) were selected for fragmentation and dynamically excluded for 0.4 min (exclusion window: mass width 0.015 m/z; 1/K0 width 0.015 Vs/cm^2^). The collision energy was set to 20 eV at 0.6 Vs/cm^2^ and 59 eV at 1.6 Vs/cm2. The 1/K0 values in between were interpolated linearly and kept constant above or below. The quadrupole isolation width was set to 2 m/z for 700 m/z and to 3 m/z for 800 m/z. Isolation width was constant except for linear interpolation between specified points. For calibration of the TIMS elution voltage, the Agilent ESI-Low Tuning Mix was used with three selected ions (m/z, 1/K0: 622.0290, 0.9915; 922.0098, 1.1986; 1221.9906, 1.3934). Mass calibration is performed with sodium formate in HPC mode. The MS data was acquired in diaPASEF. 10 diaPASEF scans with 2 ion mobility windows per scan were identified as ideal, resulting in an average of 7 - 8 datapoints per peak based on the average peak width determined in DDA PASEF mode (9s). py_diAID (77) was employed for the optimal positioning of dia-PASEF isolation windows. Identified was an optimal mass range of 285.9 to 1476.7 m/z and an ion mobility of 0.7 to 1.35 Vs/cm^2^.

### DIA data processing

The raw TIMS data (.d folders) were directly loaded in DIA-NN (version 1.8.1) (78). The mouse protein FASTA file (UP000000589, reviewed, 08.2020) was used in the library-free mode for a library generation with deep learning-based sepctra, RTs and IMs prediction enabled. The precursor ion generation parameters were as follows: Trypsin/P with max 1 missed cleavage, max 0 variable modifications, N-terminal M excision and C carbamidomethylation enabled, peptide length from 7 - 30, precursor charge range 2 - 4, precursor m/z range 280 – 1500, fragment ion m/z range 200 – 1800. The precursor FDR were set to 1%. The algorithm parameters were set as follows: mass accuracy, MS1 accuracy and scan window were set to 0. The use of isotopologues, MBR and no shared spectra were enabled. Protein inference on genes level, neural network classifier in single-pass mode, quantification strategy set to robust LC (high precision), cross-run normalization RT-dependent, library generation was set to smart profiling and the speed and RAM usage set to receive optimal results. The report.unique_genes_matrix output (MaxLFQ, normalized and filtered with q value <0.01) generated by DIA-NN was used for further data analysis.

### Data-analysis Proteomics

Proteins that were identified in less than two EV replicates were removed. Data was log2 transformed and imputation was performed in three steps. Proteins with one missing value were considered as missing-at-random and were imputed using the missForest package in R 4.3.1 with default settings. Proteins with two or three missing values (only in N2A samples) were considered as missing-not-at-random and imputed by randomly drawing from a gaussian distribution around the global minimum (3 missing values) or around the lowest value for each protein (2 missing values). For visualization, the ComplexHeatmap package was used after values were scaled. Fold change and statistical significance were determined in Perseus 2.0.11.0 using a two-tailed t-test with FDR-based correction with default settings.

Proteins that were enriched over 10-fold in EV samples compared to N2A cell lysates were analyzed for GO term enrichment using the DAVID functional annotation tool (accessed 17.10.2023). The protein list was tested for enrichment of GO terms for cellular compartment (GO_TERM_DIRECT) against the whole mouse proteome.

### Primary cell isolation

Animal procedures were approved by the Ethics committee for Animal Experiments and the Animal Welfare Body of Leiden University (AVD10600202215851; 15851,1-193) and were performed in accordance with the guidelines of the Dutch government and the European Directive 2010/63/EU. Wild-type C57BL/6J mice were bred in-house and kept in a temperature-controlled room with a 12 hours-12 hours light-dark cycle. Food and water were provided *ad libitum*. Primary neurons were isolated from early postnatal mouse hippocampus based on a protocol by Beaudoin et. al(79). 0-2 day old (P0-2) C57BL/6J mice were decapitated and hippocampi were dissected. Tissue was first dissociated with 0.25% trypsin at 37°C for 20 min, then DNase (0.1% (w/v), Sigma Aldrich, DN25) was added and incubated for 5 min at room temperature. Cells were triturated, counted as described above and 100.000 cells were seeded on poly-lysine coated 35 mm glass-bottom dishes (ibidi, 81158) in Neurobasal medium (Gibco) containing 2% B-27 supplement (Gibco), 1% GlutaMAX (Gibco), and penicillin-streptomycin (200 µg/mL each), and grown at 37°C in 5% CO_2_. 2 days after plating, cytosine b-D-arabinofuranoside (AraC) (Sigma Aldrich, C6645) was added in a 50% medium change (final concentration 2 µM). After 24-48h, medium was fully exchanged to remove AraC. Then, 50% of medium was refreshed every 3-4 days until cultures were used at day in vitro 11-15.

### Microscopy primary cells

HEK293T_eCB2.0_ cells were dissociated and counted as described above. 12h prior to imaging, 80.000 cells were seeded on top of the hippocampal neurons in a 50% medium change. The following day, medium was replaced with 540 µL phenol-red free Neurobasal medium (2% B-27, 1% GlutaMAX, 200 µg/mL penicillin-streptomycin) containing 0.1% DMSO or inhibitor and incubated for 20 minutes in the incubator before placing the dish on the microscope stage. Images were acquired and processed as described above except for the following changes: A brightfield image was captured prior to acquisition using a Nikon DS-Fi3 camera. Only the 488nm laser was used for excitation. 60 µL 10x (S)-3,5-DHPG (final concentration 50 µM, Cayman Chemical, 14411) was added.

### Animals

Animal experiments conducted in the United States were approved by the Institutional Animal Care and Use Committee of Indiana University and conform to the National Institutes of Health Guidelines on the Care and Use of Animals. Mice were kept under approved, specific pathogen– free laboratory conditions (12-hour light/12-hour dark cycle, 22° to 24°C, 40 to 70% humidity), and all efforts were made to minimize pain and suffering and to reduce the number of animals used. In electrophysiological experiments, C57Bl/6 J male mice (postnatal days 27 to 45) were utilized.

### Acute slice preparation

Mice were decapitated under deep isoflurane anesthesia. The brains were carefully removed from the skull and transferred rapidly to ice-cold sucrose containing artificial cerebrospinal fluid (sucrose-ACSF; containing 75 mM NaCl, 75 mM sucrose, 2.5 mM KCl, 25 mM glucose, 1.25 mM NaH2PO4, 4 mM MgCl2, 0.5 mM CaCl, and 24 mM NaHCO3, Sigma-Aldrich, St. Louis, MO, USA), equilibrated with 95% O2 and 5% CO2. All chemicals and reagents were purchased from Sigma-Aldrich, unless mentioned otherwise. Coronal hippocampal acute slices (300 μm thick) were cut with a VT-1200S Vibratome (Leica, Nussloch, Germany) (anteroposterior −1.8 to −2.8 mm from bregma) and were incubated in sucrose-ACSF for 1 hour at 34°C. Afterward, the oxygenated incubation chamber was kept at room temperature, and slices were subjected to subsequent electrophysiological experiments.

### In vitro slice electrophysiology

All electrophysiological recordings were made in a submerged recording chamber at 33°C constantly perfused with oxygenated recording ACSF solution (126 mM NaCl, 2.5 mM KCl, 10 mM glucose, 1.25 mM NaH2PO4, 2 mM MgCl2, 2 mM CaCl2, and 26 mM NaHCO3). Slices were visualized with an upright Nikon Eclipse FN1 microscope equipped with infrared differential interference contrast (DIC) optics (Nikon, Tokyo, Japan).

Whole-cell patch-camp recordings were obtained from interneurons in the CA1 region of the hippocampus after visual inspection of their somatic location in the radiatum layer and their multipolar morphology under a DIC microscope. Recordings were carried out with borosilicate glass pipettes (0.86-mm inner diameter and 1.5-mm outer diameter with 3- to 5-megohm resistance) filled with internal solution (containing 126 mM K-gluconate, 4 mM KCl, 10 mM Hepes, 4 mM Mg– adenosine triphosphate (ATP), 0.3 mM Na2–guanosine triphosphate (GTP), 10 mM phosphocreatine, and 8 mM biocytin; pH 7.2; 290 mOsm/kg). Pipettes were pulled with a P-1000 horizontal micropipette puller (Sutter Instrument, Novato, CA, USA). For paired recordings, pyramidal cells were selected in the pyramidal layer of the hippocampal CA1 region and were recorded in whole-cell voltage-clamp configuration (holding potential was set to −70 mV). Postsynaptic internal solution (containing the following: 40 mM CsCl, 90 mM K-gluconate, 1.8 mM NaCl, 1.7 mM MgCl2, 3.5 mM KCl, 0.05 mM EGTA, 10 mM Hepes, 2 mM Mg-ATP, 0.4 mM Na2-GTP, and 10 mM phosphocreatine; pH 7.2; 290 mOsm/kg) was also supplemented with either SecinH3 (10 µM) and NAV2729 (30 µM), WOBE437 (10 µM) or with 0.5% vehicle DMSO for control. Throughout paired recordings, series resistances were carefully monitored, and recordings were discarded if the series resistance changed >20% or reached 25 megaohms. Action potentials in presynaptic interneurons were elicited in current-clamp mode by injecting 2-ms-long 2-nA square pulses at 10-Hz frequency. DSI was induced in every 2 minutes throughout the recording by using a 1-s-long depolarization pulse on the pyramidal cell from −70 to 0 mV. IPSC charges were then analysed and compared between the pre-DSI period and the post-DSI period. The charge of an IPSC is calculated as the integral of the current over the time course of the synaptic event. Recordings were performed using MultiClamp 700B amplifiers (Molecular Devices, San José, CA, USA). Signals were filtered at 3 kHz using a Bessel filter and digitized at 10 kHz with Digidata 1550 analog-to-digital interface (Molecular Devices). The recorded traces were analyzed using the Clampfit 10 software (Molecular Devices).

### HEK239T_eCB2.0_ membrane preparation

HEK239T_eCB2.0_ cells were cultured in 15-cm diameter plates and were collected by scraping in 5 mL phosphate-buffered saline (PBS) and centrifuged at 5,000 x g for 5 min. Pellets derived from 30 plates were combined and resuspended in 20 mL cold Tris-HCl, MgCl_2_ buffer (50 mM Tris-HCl (pH 7.4), 5 mM MgCl_2_). The cell suspension was homogenized using an UltraTurrax homogenizer (Heidolph Instruments Schwabach, Germany). Membranes and cytosolic fractions were separated by centrifugation in a Beckman Optima LE-80K ultracentrifuge (Beckman Coulter Inc., Fullerton, CA, USA) at 100,000 g for 20 min at 4°C. The supernatant was discarded. The pellet was resuspended in 10 mL cold Tris-HCl, MgCl_2_ buffer and homogenization and centrifugation steps were repeated. The membranes were resuspended in 10 mL cold Tris-HCl, MgCl_2_ buffer. Aliquots of 100 µL were stored at -70°C until further use. The protein concentration was determined using the Pierce™ BCA Protein Assay Kit (ThermoFisher Scientific, Waltham, MA, USA).

### Homologous and heterologous [^3^H]CP-55,940 displacement assays

HEK293T_eCB2.0_ membrane aliquots of 20 µg membrane protein were used for the displacement assays. For assays with endocannabinoids, the membranes were preincubated for 30 min with 50 µM PMSF in assay buffer (50 mM Tris-HCl (pH 7.4), 5 mM MgCl_2_, 0.1% BSA). The homologous displacement assays were performed with a range of concentrations CP-55,940 in the presence of either ∼3 nM, ∼6 nM, or ∼18 nM [^3^H]CP-55,940 (specific activity 106.5 Ci/mmol; Revvity, Waltham, MA) in assay buffer. The heterologous displacement assays were performed with a range of endocannabinoid concentrations in the presence of ∼13 nM [^3^H]CP-55,940 in assay buffer. For both assays, binding was initiated by addition of membrane homogenates to reach a final volume of 100 µL. Total binding was determined in absence of competing ligand. Nonspecific binding was determined in the presence of 10 µM Rimonabant. The final concentration of organic solvent (DMSO for Rimonabant or acetonitrile for endocannabinoids) was 0.25% (v/v) in assay buffer in all samples. Total radioligand binding did not exceed 10% of the amount added to prevent ligand depletion. Incubation was done at 25°C for 2h to reach equilibrium. Incubations were terminated by rapid vacuum filtration with ice-cold assay buffer through GF/C 96-well filter plates (Revvity, Waltham, MA using a Filtermate-96-well harvester (Revvity, Groningen, The Netherlands). Filters were subsequently washed twenty times with ice-cold assay buffer. Filters were dried for at least 30 min at 55°C, after which 25 μl MicroScint-O (Revvity, Groningen, The Netherlands) was added per well, followed by a 3h incubation. The filter-bound radioactivity was determined by scintillation spectrometry using a Microbeta2® 2450 microplate counter (Revvity, Boston, MA).

### Mathematical model of signal formation dynamics

A semi-mechanistic mathematical model based on ordinary differential equations was developed to capture the signal formation dynamics of the Neuro2a-GRAB 2-AG signalling system (Supporting Information). The model included the production, metabolism, and accumulation of 2-AG in Neuro2A cells, as well as its release into the extracellular space, and was fitted to the available time course fluorescence signal, while fixing several separately determined model parameters. Mixed-effects modeling was used to account for variability in fluorescence signal between wells associated with cell densities. Model fitting and simulations were performed using R (version 4.2.1).

### Statistical Analysis

Statistical analysis of proteomics data was performed as described above. For all other data statistical analysis was done in GraphPad Prism 8.4.3 or 9.0.0 as stated in the respective figure legends.

## Author contributions

V.M.S and M.v.d.S. conceived the project. V.M.S, B.B., S.T.T, C.v.d.H., N.S., I.K. and M.v.d.S. designed the experiments. V.M.S performed the majority of experiments. B.B. performed electrophysiology experiments. S.T.T performed mathematical modeling. J.R. and A.F.S performed mass spectrometry measurements. C.v.d.H. performed radioligand binding assays. V.M.S, B.B., S.T.T., C.v.d.H., A.F.S., J.R., C.v.d.H., L.H.H., J.G.C.v.H., Y.L., N.S., I.K. and M.v.d.S. analyzed and interpreted the data. V.M.S and M.v.d.S wrote the manuscript with input from all authors.

## Acknowledgements

We thank the Netherlands Organization for Scientific Research VICI-grant, 724.017.002 (MvdS).

