## Supplementary Information for "The endocannabinoid 2-arachidonoylglycerol is released and transported on demand via extracellular microvesicles"

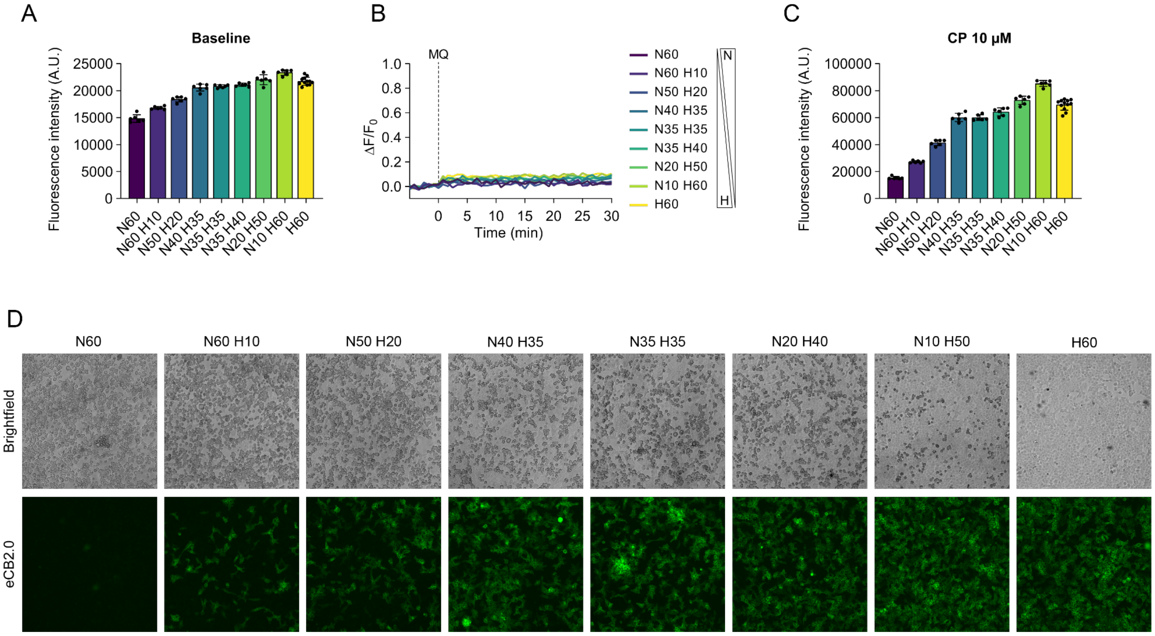


**Supplementary Figure 1. Optimization of the endocannabinoid transport assay. (A)** Absolute baseline fluorescence of different number of $HEK293T_{eCB2.0}$ and Neuro2A cells. **(B)** Changes in fluorescence compared to baseline (ΔF/F0) after treatment with vehicle (MQ). **(C)** Absolute fluorescence after addition of $CB_1$ receptor agonist (-)CP-55,940 (10 µM). Data in A-C are mean of 3-6 wells in a 96-well plate. Error bars in A and C represent SD. **(D)** Brightfield and fluorescence widefield images of selected wells of each cell combination after addition of (-)CP-55,940 (C). Round cells with high contrast in the brightfield image are Neuro2A cells. Stretched out cells are $HEK293T_{eCB2.0}$ cells and are visible in the fluorescent image. Images were acquired with an EVOS® FL Auto Imaging System.

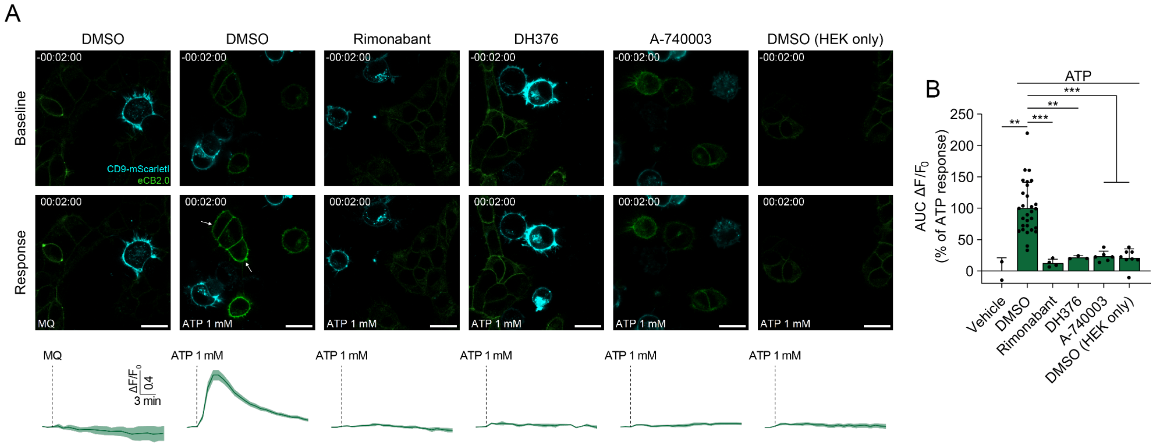


**Supplementary Figure 2. Live-cell confocal microscopy illuminates basic mechanisms of 2-AG signaling. (A)** Representative images of HEK293T$_{eCB2.0}$ cells and Neuro2A cells transiently expressing CD9-mScarletI before (Baseline, 3 min) and during (Response, 20 min) ATP stimulation after treatment with DMSO, 1 µM Rimonabant, 1 µM DH376 or 10 µM A-740003 for 20 min. Traces show mean fluorescence changes ($\Delta F/F_0 \pm$SEM) of 2-28 ROIs from 2-15 individual dishes. Scale bars are 20 µm. **(B)** AUC of traces in (A); percentage of vehicle-corrected ATP-response. Data are shown as mean±SD. Statistical significance was determined by one-way ANOVA with Tukey's correction for multiple comparisons. ** $P < 0.01$, *** $P < 0.001$.

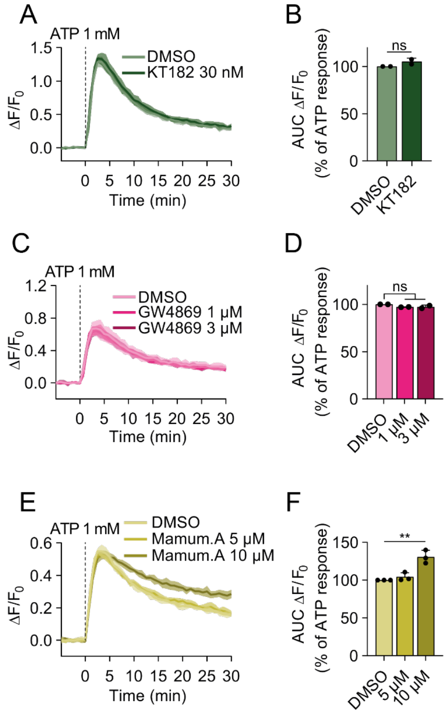


**Supplementary Figure 3. ABHD6 inhibition and exosome release inhibitors do not block 2-AG release. (A,C,E)** Representative traces (mean±SD; N=1, n=6) of changes in fluorescence upon ATP-stimulation in the endocannabinoid transport assay. Cells were treated with the indicated compounds for 30 min at 37°C prior to baseline measurement. **(B,D,F)** Area under the curve (AUC) of fluorescent changes ($\Delta F/F_0$), calculated as percentage of vehicle-corrected ATP-response. Data is shown as mean±SD (N=2-3, n=6). Statistical analysis was performed using two-tailed t-test (B) or one-way ANOVA with Tukey's correction for multiple comparisons (D,F). ** $P < 0.01$, ns not significant $P > 0.05$.

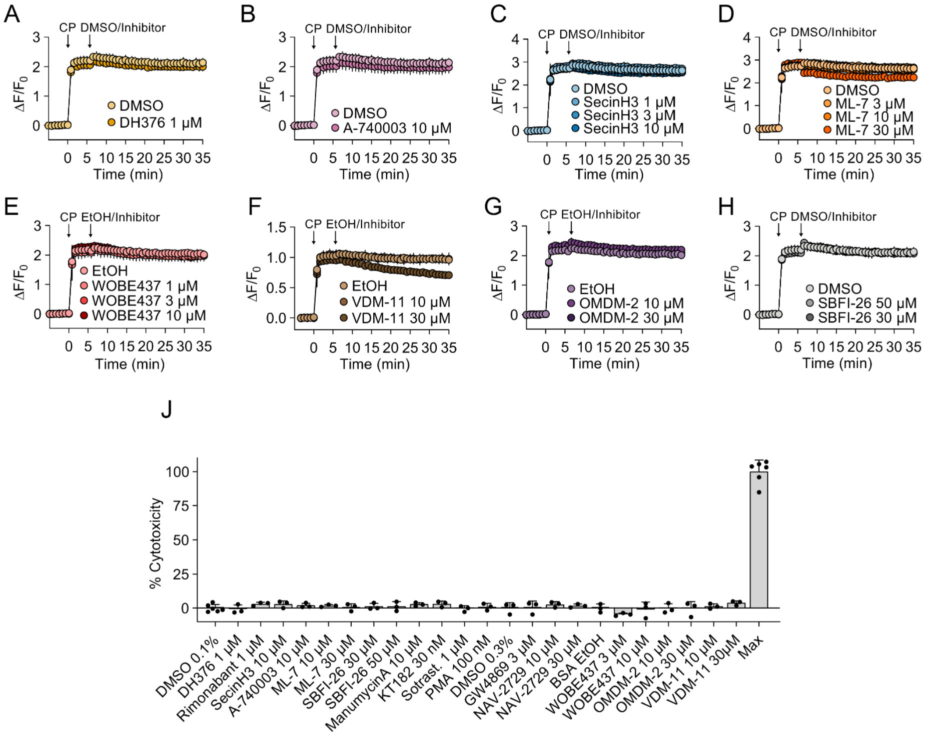


**Supplementary Figure 4. Influence on compounds on CP-activated GRAB$_{eCB2.0}$ signal in HEK293T cells. (A-H)** Changes in GRAB$_{eCB2.0}$ fluorescence ($\Delta F/F_0$) in HEK293T cells. 10 µM of (-)CP-55,940 was added after baseline measurement. After 5 minutes, compounds were added at the indicated concentrations and the effect on $\Delta F/F_0$ was monitored for 30 minutes. **(I)** Neuro2A and HEK293T$_{eCB2.0}$ cells in co-culture were treated with the indicated compounds for 1h at 37°C and LDH activity was measured in the supernatant to estimate cytotoxicity. Cytotoxicity was calculated relative to a positive control (lysis buffer). Data is represented as mean±SD (n=3-6 wells of a 96-well plate).

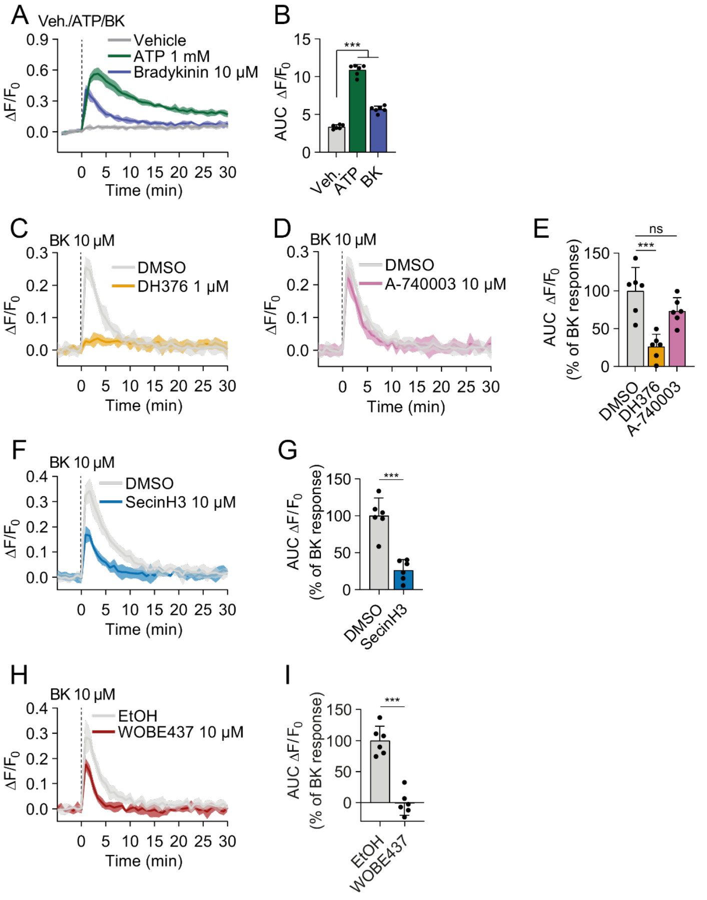


**Supplementary Figure 5. Bradykinin-stimulated 2-AG mobilization is independent of $P2X_7R$ and can be blocked by WOBE437 and SecinH3. (A,C,D,F,H)** Traces of changes in fluorescence upon ATP- or BK-stimulation in the co-culture endocannabinoid transport assay. Cells were treated with the indicated compounds for 30 min at 37°C prior to baseline measurement. **(B,E,G,I)** Area under the curve (AUC) of fluorescent changes ($\Delta F/F_0$) in (A,C,D,F,H). All data are shown as mean±SD (N=1, n=6). Statistical analysis was performed using one-way ANOVA with Tukey's correction for multiple comparisons. *** $P < 0.001$, ns not significant $P > 0.05$.

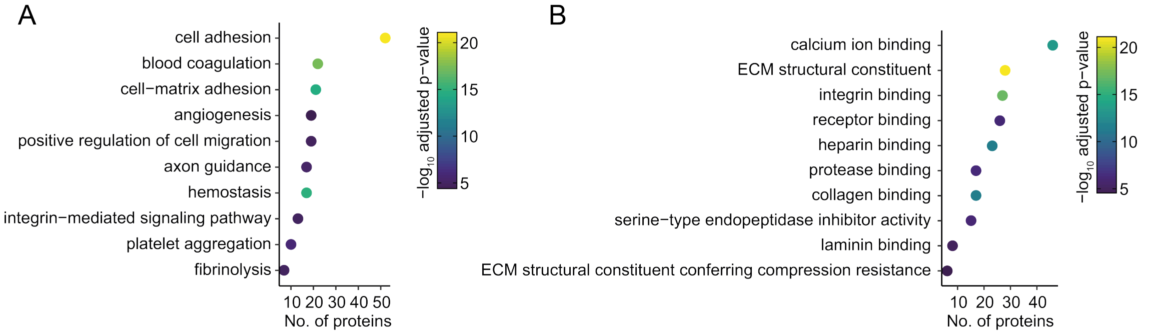


**Supplementary Figure 6. Molecular function and biological process gene ontology of EV proteome (A,B)** Gene ontology enrichment analysis for biological process (GO_BP_DIRECT, A) and molecular function (GO_MF_DIRECT, B) of proteins enriched >10-fold in EVs compared to Neuro2A cell lysate as determined by LC-MS/MS based proteomics.

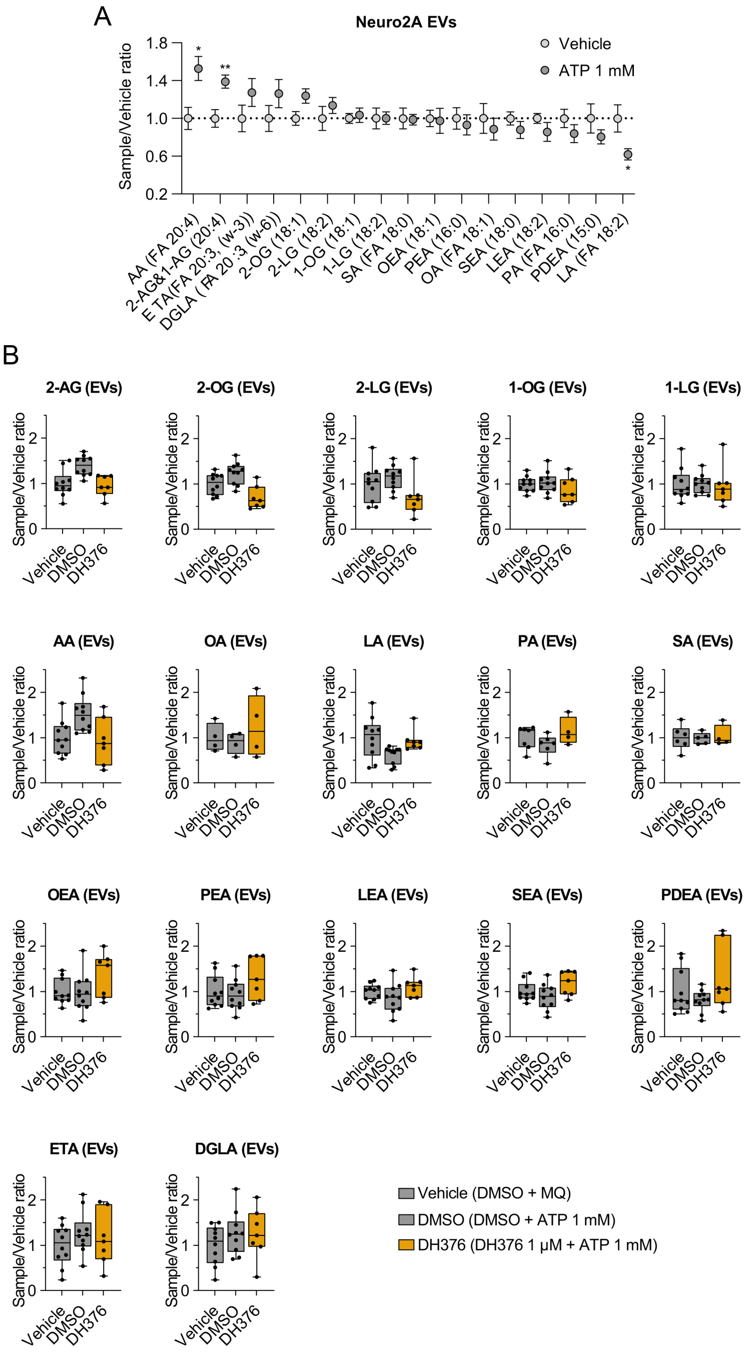


**Supplementary Figure 7. Lipids identified in EVs by targeted LC-MS/MS based lipidomics. (A)** Fold-enrichment (ATP/Vehicle) of lipids in EVs following vehicle (MQ) or ATP-treatment for 30 min at 37°C. Data are shown as mean±SEM. Statistical significance was determined using one-way ANOVA with Tukey's correction for multiple comparisons. * $P$ < 0.05, ** $P$ < 0.01, ns not significant $P$ > 0.05. N=7-10 EV isolations. **(B)** Lipid levels in EVs relative to vehicle-treated control. Cells were treated with 1 µM DH376 or DMSO for 20 min prior to vehicle (MQ) or ATP-treatment for 30 min at 37°C, followed by EV isolation.

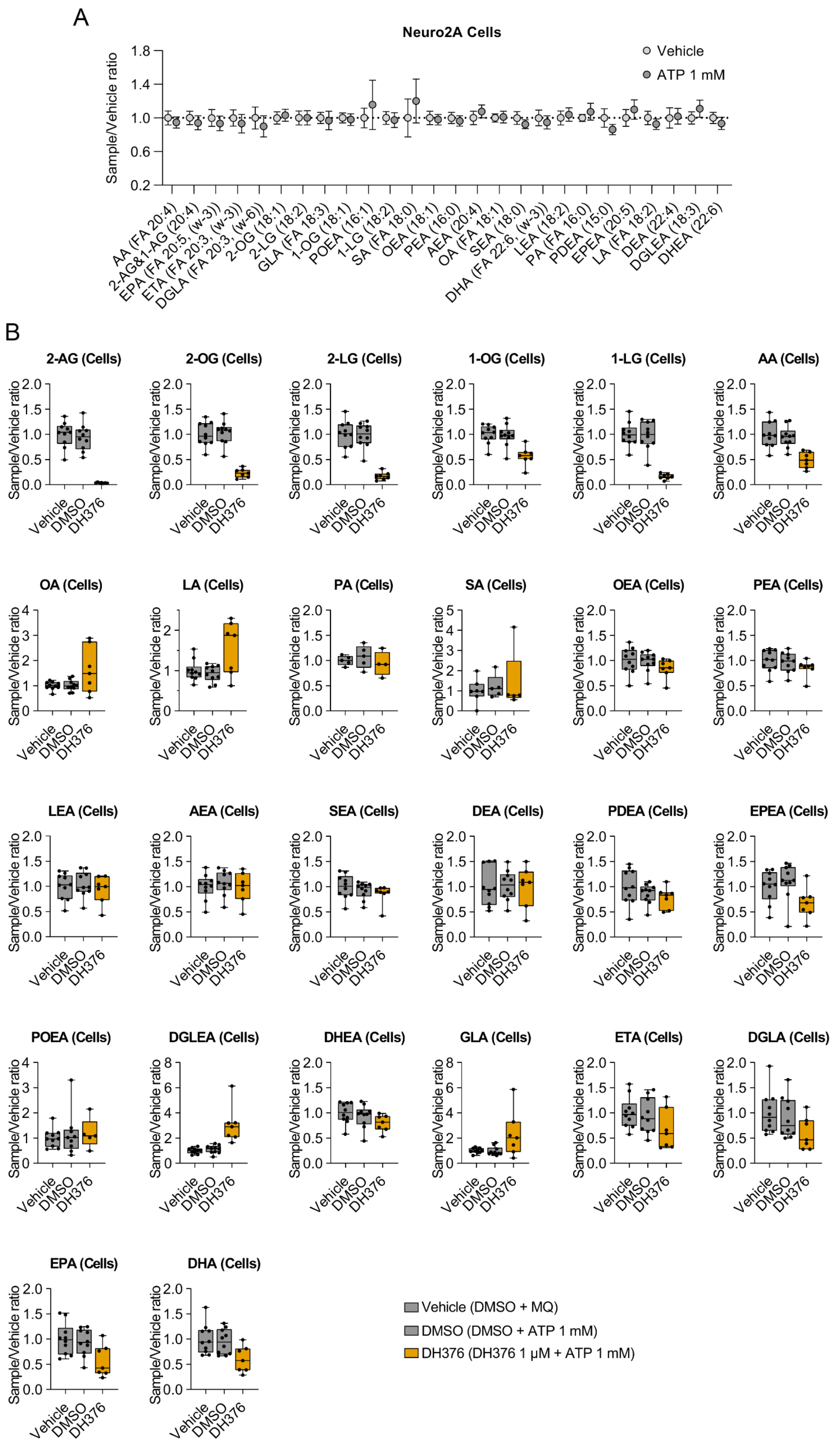


**Supplementary Figure 8. Full lipidomics cells. (A)** Fold-enrichment (ATP/Vehicle) of lipids in cells following vehicle (MQ) or ATP-treatment for 30 min at 37°C. Data are shown as mean±SEM. **(B)** Lipid levels in cells relative to vehicle-treated control. Cells were treated with 1 μM DH376 or DMSO for 20 min prior to vehicle (MQ) or ATP-treatment for 30 min at 37°C. N=7-10.

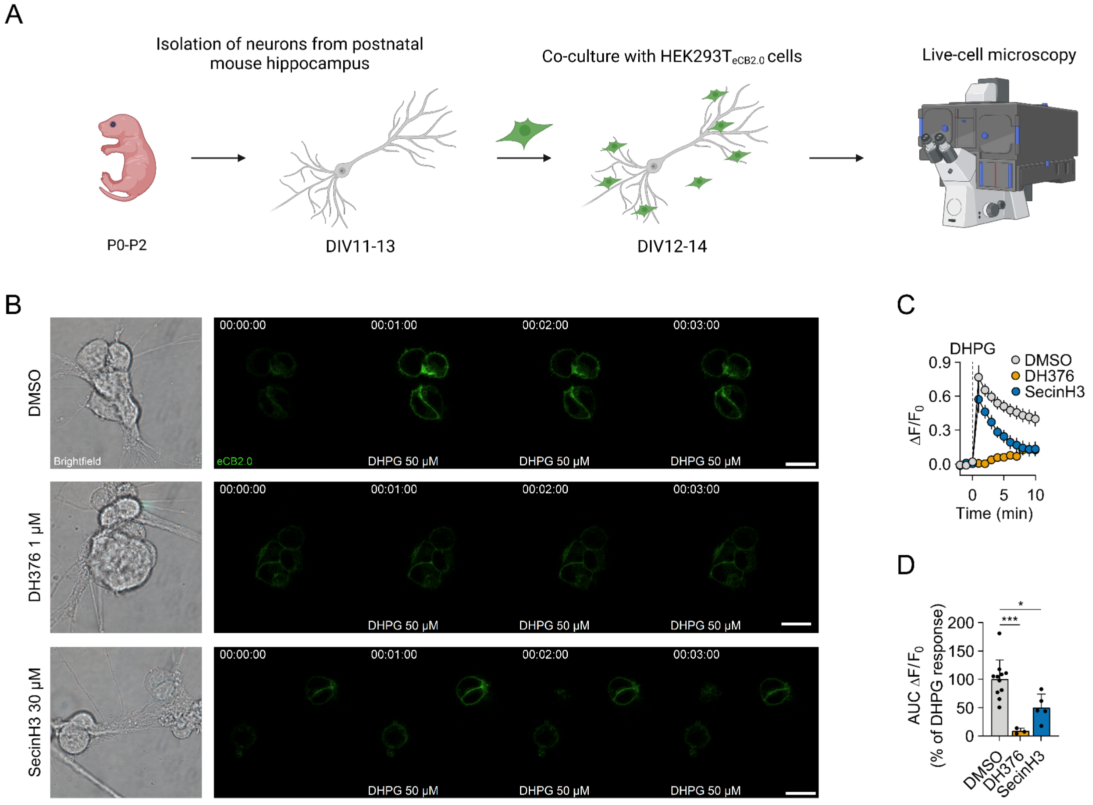


**Supplementary Figure 9. Arf6 and DAGL control 2-AG mobilization in hippocampal neurons. (A)** Hippocampal neurons were isolated from early postnatal mouse brain and matured until day in vitro (DIV) 11-13. HEK293T$_{eCB2.0}$ cells were then seeded on top and GRAB$_{eCB2.0}$ activation was recorded the following day using confocal live-cell microscopy. **(B)** Representative brightfield and confocal images of HEK293T$_{eCB2.0}$ cells and mouse hippocampal neurons before and after stimulation with 50 µM DHPG. Cells were treated with the indicated compounds for 20 min prior to addition of DHPG. Scale bars are 20 µm. **(C)** Traces show mean fluorescence changes ($\Delta F/F_0$±SEM) of 3-11 ROIs from 2-5 individual dishes. **(D)** AUC of traces in (C) calculated as percentage of vehicle-corrected DHPG-response. Data are mean±SD. Statistical significance was determined by one-way ANOVA with Tukey's correction for multiple comparisons. * $P < 0.05$, *** $P < 0.001$.

**Modeling Objective and Summary**

Mathematical modeling was performed to determine the rate-limiting step in 2-AG release in the co-culture transport assay. Two models were developed. Both models assumed a constant production and metabolism of 2-AG in Neuro2A. The release of 2-AG from Neuro2A cells to the extracellular space was assumed to be dependent on ATP stimulus. The first model assumed the release of 2-AG in absence of accumulated 2-AG pool in Neuro2A cells upon an ATP stimulus. The second model assumed the presence of a pool of 2-AG molecules in the forming extracellular vesicles before they are released from Neuro2A cells. An ATP stimulus rapidly released the EVs. Both models were evaluated based on its ability to describe the observed fluorescence signal kinetics produced by $HEK_{293T.GRABeCB2.0}$ cells upon 2-AG uptake. Additional experimental information was used to derive the steady-state diacylglycerol (DAG) levels, DAG-dependent 2-AG production rate constant, and 2-AG metabolism rate constant in Neuro2A cells.

**Methods**

***Model Structure***

A. **Model Assumptions**

Models used in the current analysis assumed the following system characteristics.

1) The amount of DAG in N2A cells was approximately constant ($m_{DAG}$≈constant).
2) In the absence of ATP, 2-AG export/release from Neuro2A cell is negligible.
3) In the presence of ATP, the rate of 2-AG release from the 2-AG EV pool ($\boldsymbol{m_{EV}}$) is rapid with respect to its production, metabolism, and distribution in the Neuro2A cell ($k_{release}$ >> $k_{production}$, $k_{metabolism,N2A}$, $k_{fill}$, $k_{sink}$). This assumption is relevant only to the pre-stimulus accumulation model.
4) In the experiment, steady state was reached before ATP was added to the environment.
5) The size of cellular 2-AG pool in Neuro2A cell is constant ($m_{cell}$≈$m_{cell,SS}$ is constant).
6) Experimentally derived parameter values were measured by assuming the presence of 40,000 N2A cells ($N_{N2A}$) and 35,000 HEK ($N_{HEK}$) cells, where applicable.
   - The exact number of Neuro2a cells for each set of observations were allowed to vary as inter-occasional variability.
   - No cell growth and death events occurred during each observation.
7) Extracellular 2-AG is partially removed from the system with a first order rate constant $k_{removal}$ due to a combination of uptake by non-signal-producing cells, degradation, and distribution to the environment, making these molecules unattainable by HEK cells.
8) The volume of HEK cell lipid bilayer plasma membrane ($V_{HEK}$) was approximately 2.65 x $10^{-15}$ L.

B. **Production-only model**

The production-only model assumed that, upon receiving ATP stimulus, 2-AG was exported from Neuro2A at a rate that is proportional to the size of cellular 2-AG pool ($m_{cell}$) in Neuro2A cells (Fig. A1a).

The size of the cellular 2-AG pool ($m_{cell}$) before and after ATP stimulus remained relatively constant. Thus, in this model, the mass balance of 2-AG in the extracellular space and HEK cells were given as the following (Eqs. A1-2).

$$\frac{dm_{extracellular}}{dt} = k_{export}N_{N2A}m_{cell} - \left(k_{removal} + k_{absorption}N_{HEK}\right)m_{extracellular} \quad \text{Eq. A1}$$

$$\frac{dm_{HEK}}{dt} = k_{absorption}m_{extracellular} - k_{metabolsim,HEK}m_{HEK} \quad \text{Eq. A2}$$

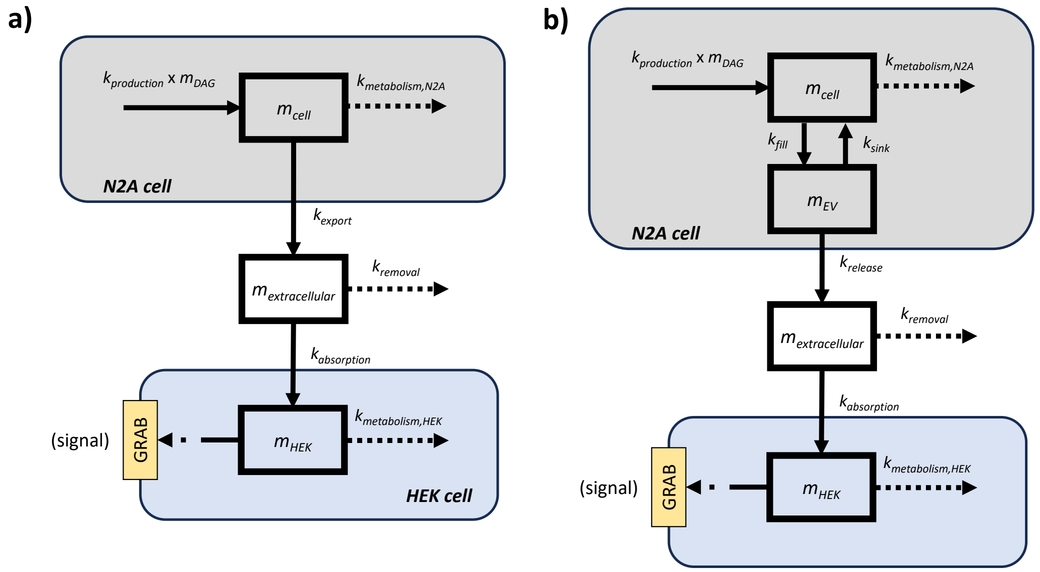


**Fig. A1 | Model illustration.** 2-AG is produced and released by Neuro2A cells after ATP stimulation and absorbed by HEK cells. GRAB sensor activity in HEK cells is dependent on 2-AG levels in the cell membrane. The production-only model **a)** assumed that 2-AG is produced and released by Neuro2A cells without accumulation. The pre-stimulus accumulation model **b)** introduced an EV pool in which 2-AG accumulates in Neuro2A cells.

The rate constant of 2-AG release ($k_{release}$) was calculable from the steady-state size of the cellular 2-AG pool ($m_{cell,SS}$; 8.85 x $10^{-17}$ mol) and the total extracellular 2-AG amount observed 30 minutes after ATP stimulus in the absence of cells that uptake 2-AG ($m_{dose}$; Eq. A3).

$$k_{export} = \frac{m_{dose}}{m_{cell,SS} \times \Delta t} = 1.39 \times 10^{-6}\ s^{-1} \qquad \text{Eq. A3}$$

Model fitting of the production-only model was performed to estimate the values of $k_{removal}$, $k_{absorption}$, and $k_{metabolism,HEK}$.

#### C. Pre-stimulus accumulation model

The current model assumed the presence of 2-AG in extracellular vesicles (EV pool), before they are released from the cell, which reached an equilibrium with the cellular 2-AG pool in Neuro2A cells before receiving ATP stimulus (Fig. A1b). This equilibrium was described using the first order rate constants $k_{fill}$ and $k_{sink}$. Here, EV pool fill rate represents the combined rate of EV formation and 2-AG loading into the EVs, while the sink rate represents the opposite phenomenon. In the presence of ATP, the export of 2-AG from the cellular pool to the extracellular space thus became a two-step process, involving the EV pool fill rate and 2-AG release rate from the EV pool. In the absence of ATP, the mass balance of 2-AG in the EV pool is expressed as the following.

$$\frac{dm_{EV}}{dt} = k_{fill} \times m_{cell} - k_{sink} \times m_{EV} \qquad \text{Eq. A4}$$

Here, the rate of 2-AG release ($k_{release}$) was assumed to be rapid with respect to $k_{fill}$ and $k_{sink}$. Therefore, in in the presence of ATP, the effective rate of 2-AG export from the

Neuro2A cell is determined by EV pool fill rate (Eq. A5). 2-AG mass balance inside the HEK cell is given by Eq. A2.

$$\frac{dm_{extracellular}}{dt} = k_{fill} N_{N2A} m_{cell} - \left(k_{removal} + k_{absorption} N_{HEK}\right) \times m_{extracellular} \quad \text{Eq. A5}$$

Model fitting of the production-only model was performed to estimate the values of $k_{removal}$, $k_{absorption}$, $k_{metabolism,HEK}$, and the pre-stimulus steady-state 2-AG amount accumulated in the EV pool ($\boldsymbol{m_{EV,SS}}$). The fill ($k_{fill}$) and sink ($k_{sink}$) rate constants of the EV pool were calculated as a function of the model-estimated value of $\boldsymbol{m_{EV,SS}}$ and by using experimentally determined values of $m_{cell,SS}$ and $m_{dose}$.

$$k_{fill} = \frac{m_{dose} - m_{EV,SS}}{m_{cell,SS} \times \Delta t} \quad \text{Eq. A6}$$

$$k_{sink} = \frac{k_{fill} \times m_{pool,SS}}{m_{signal,SS}} \quad \text{Eq. A7}$$

#### *Experimentally Determined Parameters*

##### A. Signal generation by GRAB sensor

GRAB sensor generates fluorescence signal upon binding to intracellular 2-AG in HEK cells. Signal generation kinetics for each HEK cell was given by the following equation.

$$S_{observed} = S_{max,HEK} \frac{C_{HEK}}{C_{2AG,HEK} + K_i} \times N_{HEK} \quad \text{Eq. A8}$$

In the above equation, $S_{observed}$ represents the observed signal level. $S_{max,HEK}$ represents the maximum signal emitted by a single HEK cell (6.29 x $10^{-5}$ ;a.u.). The signal was dependent on intracellular HEK 2-AG concentration ($C_{HEK}$), which is calculated using a plasma membrane volume $V_{HEK}$ of approximately 2.65 x $10^{-15}$ L. The total number of HEK cells involved in each set of observation was given by $N_{HEK}$.

$K_i$ represents the binding constant of 2-AG to the GRAB sensors (4.17 x $10^{-5}$ mol/L) determined as described in the material and methods. Notably, we observed an approximately 15-fold lower binding affinity of the $GRAB_{eCB2.0}$ sensor for 2-AG compared to wildtype $CB_1R$. This has two implications. First, it is unlikely that the sensor acts as a sink, reducing the availability of 2-AG molecules for the $CB_1R$. Second, the sensor underestimates the physiological activation of the $CB_1R$ in the brain, because less 2-AG molecules are needed to achieve full receptor occupancy and to activate the wildtype receptor. This should be considered, when analyzing $GRAB_{eCB2.0}$ sensor data.

##### B. 2-AG metabolism and production rate constant in Neuro2A cells

The metabolism rate constant of 2-AG in Neuro2A cells ($k_{metabolism,N2A}$) was derived from the measured steady-state 2-AG amount in the cellular N2A pool ($m_{cell,SS}$=8.85 x $10^{-17}$ mol) and its amount 30 minutes after 2-AG production was artificially inhibited ($m_{cell,inhibited}(30\ minutes)$=1.84 x $10^{-18}$ mol). Assuming $k_{fill}$ and $k_{sink}$ to be sufficiently small with respect to 2-AG production and metabolism, the rate of 2-AG metabolism in the cellular Neuro2A pool is given as the following.

$$k_{metabolism,N2A} = \frac{1}{t} \ln\left(\frac{m_{cell,SS}}{m_{cell,inhibited}(30 minutes)}\right) = 2.15 \times 10^{-3}\ s^{-1} \quad \text{Eq. A9}$$

Consquently, 2-AG production rate constant is calculable ($m_{DAG}$ = 2.82 x $10^{-17}$ mol; constant).

$$k_{production} = \frac{k_{metabolism} \times m_{cell,SS}}{m_{DAG}} = 6.74 \times 10^{-3}\ s^{-1}$$ Eq. A10

#### *Parameter Fitting*

The current analysis was performed using mixed effect modeling. Mixed-effect modeling has three main structures: 1) fixed-effect (structural description of the system dynamics), 2) random-effect (statistical description of parameter variability expected between occasion), and 3) residual effect (statistical description to capture residual unexplained variability). This technique was used instead of general fixed-effect modeling method to quantify inter-occasional variability that may arise between experiments, thereby assist distinguishing the dynamic characteristics of the system from inter-occasional variability. In this analysis, inter-occasional variability was applied on the number of Neuro2a cells ($N_{N2A}$). Variability and was assumed to be log-normally distributed with random effect parameter $\eta_i$ for each occasion ***i***.

$$N_{HEK,applied,i} = N_{HEK} \times e^{\eta_i}$$ Eq. A11

Since the distribution of observations along the y-axis appeared to be independent of the mean signal level (Fig. 3), additive effect model was selected to describe the residual error of the model.

$$Prediction = Observation + \varepsilon_{residual}$$ Eq. A12

Model implementation and parameter optimization was performed on R (v. 4.2.1). Particle swarm optimization (implemented in the *pso* package) was used for initial approximation of fixed parameter value. Parameter fitting for the final mixed-effect model was performed using the FOCEi algorithm implemented in the *nlmixr2* package. Goodness of model fit was evaluated by visual predictive check, Akaike information criterion (AIC), and prediction-observation plots.

### Results

#### *2-AG accumulation was essential to describe the observed signal dynamics*

Applying the production-only model to the observed data resulted in a suboptimal model fit (AIC of -690.9; Fig. A2; Table A1). This was primarily due to the inability of the production-only model to capture the dynamics of signal peak formation observed at approximately 220 s.

**Table A1. Parameter estimates for the production-only model**

| Parameter | Description | Estimate (95% CI) | %BSV | %RSE |
|---|---|---|---|---|
| $k_{removal}$ ($s^{-1}$) | Rate of extracellular 2-AG removal | $5.22 \times 10^{-15}$ ($2.01 \times 10^{-21}$ ~ $1.36 \times 10^{-8}$) | | 22.9 |
| $k_{absorption}$ ($s^{-1}$) | Rate of 2-AG absorption into HEK cell | $4.28 \times 10^{-6}$ ($3.90$ ~ $4.71) \times 10^{-6}$ | | 0.389 |
| $k_{metabolism,HEK}$ ($s^{-1}$) | Rate of 2-AG metabolism in HEK cell | $2.07 \times 10^{-2}$ ($1.96$ ~ $2.19) \times 10^{-2}$ | | 0.73 |
| $N_{N2A}$ (count) | Number of Neuro2a cells in the experiment | $4.0 \times 10^{4}$ (fixed) | 80.5 | |
| *Residual error (additive)* | | 0.171 | | |

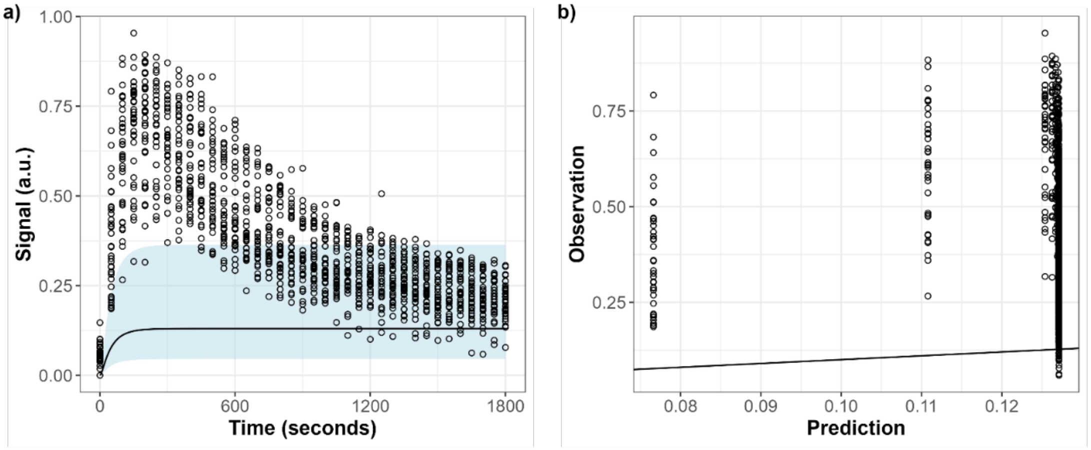


**Fig. A2 | Fit of the production-only model.** Model fitting was performed using the production-only model. **a)** Signal time-course profile observed in the experiment (points) compared to the signal profile predicted by the model (solid line). Shaded area represented the 95% prediction interval of the model fit. **b)** Observation-prediction points deviate unequally along the line of unity, suggesting an empirically inadequate model structure.

The implementation of EV pool as a source of initial 2-AG burst in the pre-stimulus accumulation model could describe the peak formation dynamics (Fig. A3, Table A2). The additional one degree of freedom used in this model was justifiable as it resulted in a statistically significant improvement of the model fit, as indicated by the drop in AIC value to -3704.2.

**Table A2. Parameter estimates for the pre-stimulus accumulation model**

| Parameter | Description | Estimate (95% CI) | %BSV | %RSE |
|---|---|---|---|---|
| $k_{removal}$ ($s^{-1}$) | Rate of extracellular 2-AG removal | $4.01 \times 10^{-3}$ $(3.05 \sim 5.28) \times 10^{-3}$ | | 2.54 |
| $k_{absorption}$ ($s^{-1}$) | Rate of 2-AG absorption into HEK cell | $1.27 \times 10^{-7}$ $(1.07 \sim 1.50) \times 10^{-7}$ | | 0.541 |
| $k_{metabolism,HEK}$ ($s^{-1}$) | Rate of 2-AG metabolism in HEK cell | $2.70 \times 10^{-3}$ $(2.42 \sim 3.00) \times 10^{-3}$ | | 0.932 |
| $m_{EV,SS}$ (mol) | Steady-state 2-AG amount the EV pool | $1.30 \times 10^{-19}$ $(1.24 \sim 1.36) \times 10^{-19}$ | | 4.43 |
| $N_{N2A}$ (count) | Number of Neuro2a cells in the experiment | $4.0 \times 10^{4}$ (fixed) | 26.8 | |
| *Residual error (additive)* | | 0.0543 | | |

The implementation of 2-AG EV pool in the model was found to significantly improve model fit. This result suggested the importance of 2-AG accumulation to describe the observed signal kinetics, especially the formation of the signal peak. This accumulation was estimated to account for approximately 58.7% of the total 2-AG dose released by Neuro2A cells over the 30 minutes observation period. Consequently, the estimated fill and sink rate constants of the EV pool were $5.73 \times 10^{-7}$ $s^{-1}$ and $3.90 \times 10^{-4}$ $s^{-1}$, respectively. These values were orders of magnitude smaller than both the production and metabolism rate constants of 2-AG in N2A cell, therefore supporting the assumption that the contribution of 2-AG distribution to the EV pool towards 2-AG mass balance in the cellular N2A pool to be negligible throughout the observation period.

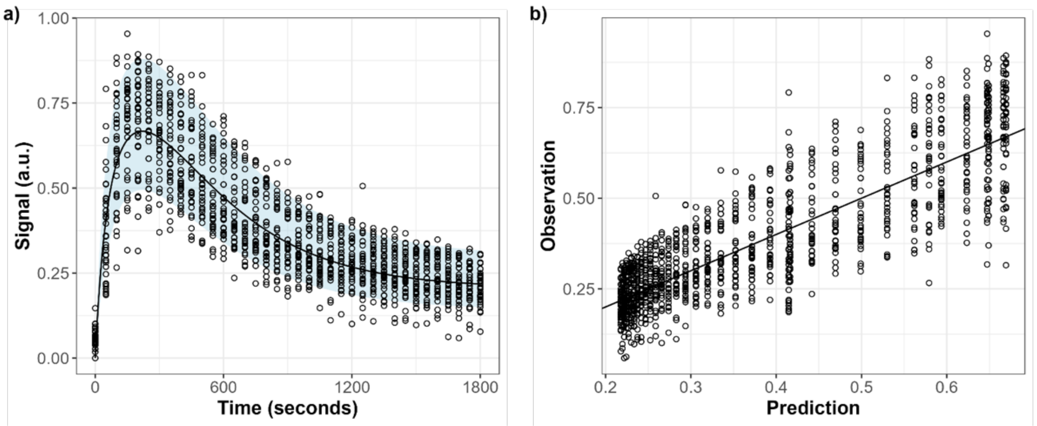


**Fig. A3 | Fit of the pre-stimulus accumulation model.** Model fitting was performed using the pre-stimulus accumulation model. **a)** Signal time-course profile observed in the experiment (points) compared to the signal profile predicted by the model (solid line). Shaded area represented the 95% prediction interval of the model fit. **b)** Observation-prediction points was distributed along the line of unity, indicating an empirically adequate model structure.

Simulation using the pre-stimulus accumulation model revealed how Neuro2A stimulation when the EV pool is empty could still lead to an increase in signal level despite not being able to create the peak signal (Fig. A4). This further supported the relevance of 2-AG accumulation in describing the formation of the signal peak.

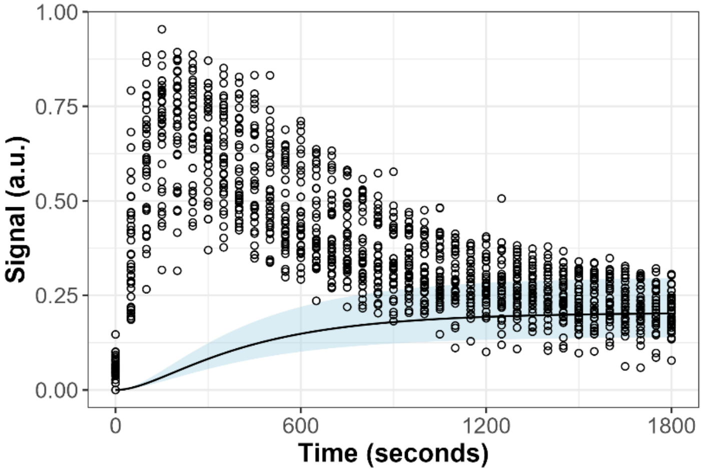


**Fig. A4 | Observed signal peak could not be reproduced without 2-AG accumulation in N2A cell.** Simulation was performed using the pre-stimulus accumulation model and by modifying model parameter $m_{EV,SS}$ to 0.

The estimated 2-AG metabolism rate constant in HEK cells 2.69 x $10^{-3}$ $s^{-1}$, equivalent to a volume-adjusted clearance ($CL_{HEK} = k_{metabolism,HEK} \times V_{HEK}$) of 7.12 x $10^{-18}$ L/s. This value was comparable to the experimentally determined 2-AG metabolism rate constant in Neuro2A cells (2.15 x $10^{-3}$ $s^{-1}$).

Each HEK cells were estimated to absorb a total of 37.6 vesicles within the 30 minutes period (assuming 2,048 2-AG molecules per-vesicle). Considering the total number of Neuro2A and HEK cells, the fraction of absorbed 2-AG relative to the total 2-AG dose was estimated to be 57.9%.

The pre-stimulus accumulation model assumes a rapid release of 2-AG from the signaling pool upon receiving ATP stimulus. Consequently, when the stimulus is introduced, the rate of vesicle release into the environment is significantly higher than it is after the initial burst of 2-AG release. Assuming a 2-AG density of 2,048 molecules per vesicle, the model estimates that approximately 38 vesicles were released at t=0, with a subsequent release rate of 0.015 vesicles/s (1 vesicle every ~67 seconds). Notably, this calculation assumed a constant 2-AG density per vesicle.
